# B cell-intrinsic STAT3-mediated support of latency and interferon suppression during murine gammaherpesvirus 68 infection revealed through an *in vivo* competition model

**DOI:** 10.1101/2023.03.22.533727

**Authors:** Chad H. Hogan, Shana M. Owens, Glennys V. Reynoso, Varvara Kirillov, Thomas J. Meyer, Monika A. Zelazowska, Bin Liu, Xiaofan Li, Aniska Chikhalya, Qiwen Dong, Camille Khairallah, Nancy C. Reich, Brian Sheridan, Kevin M. McBride, Patrick Hearing, Heather D. Hickman, J. Craig Forrest, Laurie T. Krug

## Abstract

Cancers associated with the oncogenic gammaherpesviruses, Epstein-Barr virus and Kaposi sarcoma herpesvirus, are notable for their constitutive activation of the transcription factor STAT3. To better understand the role of STAT3 during gammaherpesvirus latency and immune control, we utilized murine gammaherpesvirus 68 (MHV68) infection. Genetic deletion of STAT3 in B cells of *CD19^cre/+^Stat3^f/f^*mice reduced peak latency approximately 7-fold. However, infected *CD19^cre/+^Stat3^f/f^* mice exhibited disordered germinal centers and heightened virus-specific CD8 T cell responses compared to WT littermates. To circumvent the systemic immune alterations observed in the B cell-STAT3 knockout mice and more directly evaluate intrinsic roles for STAT3, we generated mixed bone marrow chimeras consisting of WT and STAT3-knockout B cells. Using a competitive model of infection, we discovered a dramatic reduction in latency in STAT3-knockout B cells compared to their WT B cell counterparts in the same lymphoid organ. RNA sequencing of sorted germinal center B cells revealed that STAT3 promotes proliferation and B cell processes of the germinal center but does not directly regulate viral gene expression. Last, this analysis uncovered a STAT3-dependent role for dampening type I IFN responses in newly infected B cells. Together, our data provide mechanistic insight into the role of STAT3 as a latency determinant in B cells for oncogenic gammaherpesviruses.

**IMPORTANCE:** There are no directed therapies to the latency program of the gammaherpesviruses, Epstein-Barr virus and Kaposi sarcoma herpesvirus. Activated host factor STAT3 is a hallmark of cancers caused by these viruses. We applied the murine gammaherpesvirus pathogen system to explore STAT3 function upon primary B cell infection in the host. Since STAT3 deletion in all CD19+ B cells of infected mice led to altered B and T cell responses, we generated chimeric mice with both normal and STAT3-deleted B cells. B cells lacking STAT3 failed to support virus latency compared to normal B cells from the same infected animal. Loss of STAT3 impaired B cell proliferation and differentiation and led to a striking upregulation of interferon-stimulated genes. These findings expand our understanding of STAT3-dependent processes key to its function as a pro-viral latency determinant for oncogenic gammaherpesviruses in B cells and may provide novel therapeutic targets.

## INTRODUCTION

The human gammaherpesviruses (GHVs) Epstein-Barr virus (EBV, human herpesvirus 4) and Kaposi sarcoma herpesvirus (KSHV, human herpesvirus 8) are the etiologic agents of numerous lymphomas and carcinomas that are a significant clinical burden to immune-compromised individuals, including people living with HIV. B cells are a long-term reservoir for GHV and theses viruses usurp host processes such as proliferation and differentiation to facilitate chronic infection (1-4). The identification of host factors that support latency and facilitate the emergence of GHV-driven malignancies is paramount as there are currently no treatments that directly target the latent phase of infection.

A hallmark of GHV infection and GHV cancers is the rapid and persistent activation of signal transducer and activator of transcription 3 (STAT3) (5). STAT3 is a master regulator of multiple cellular processes such as proliferation, apoptosis, and metastasis (6, 7). Constitutive STAT3 activation is reported in multiple human cancers (8, 9). Cytokines are potent activators of STAT3 in both cancer and immune cells. For instance, IL-6 induces the expression of STAT3- target genes responsible for proliferation and survival supporting tumorigenesis, while IL-10 reduces immune cell function and inflammation in some contexts (10, 11). The human GHVs use diverse strategies to promote STAT3 activation (5). EBV transmembrane protein LMP1 activates NF-κB through recruitment of cytoplasmic signaling adaptors leading to IL-6-driven STAT3 activation and cell survival (12, 13). EBV LMP2A activates STAT3 through multiple host factors to promote proliferation and survival (14, 15). The viral homologs of host cytokines, EBV vIL-10 and KSHV vIL-6, and the KSHV vGPCR also activate STAT3 (16-20).

Due to their narrow host range, the study of EBV and KSHV is largely restricted to cell culture and humanized mouse models. For that reason, murine gammaherpesvirus 68 (MHV68) is used as a model system. MHV68 is colinear with KSHV, encoding 64 direct homologs, establishes long term latency in memory B cells, and it induces lymphoproliferation in immunosuppressed mice (4). The infection of mice with MHV68 is a genetically tractable system to study virus-host interactions in specific cell types during the acute or chronic phase of infection in a natural host. Viral and host factors such as NF-κB signaling pathways support the establishment and latency in B cell compartments (4, 21-26).

The interplay of viral and host factors aids in the establishment and expansion of GHV latency, taking particular advantage of B cell processes occurring during the germinal center (GC) reaction. The GC is the site within secondary lymphoid tissues where B cells undergo maturation and differentiation processes including somatic hypermutation and class switch recombination, leading to the production of high-affinity and long-lived memory B cell and plasma cell subsets (27, 28). GHVs both engage and circumvent the GC B cell selection process to promote access to the memory B cell long-term latency reservoir for EBV and likely KSHV (29-32), and may contribute to B cell lymphomas with post-germinal center signatures (33, 34). At the peak of latency establishment ∼16 days post infection (dpi), MHV68 is predominantly found in B cells bearing GC markers (21). T follicular helper cell production of IL- 21 is required to support MHV68 latency establishment in GC B cells (23), while Blimp1 is required to access the post-GC plasma cell compartment that is a source of virus reactivation (35). The MHV68 viral M2 protein promotes host IL-10 production through the NFAT pathway, promoting proliferation and differentiation, driving B cells to a plasmablast-like phenotype *in vivo* (36-38). MHV68 RTA protein encoded by ORF50 acts synergistically with STAT3 to increase transcriptional activity in response to IL-6 (39). Given the critical role of STAT3-activating cytokines IL-10 and IL-21 in GHV infection (23, 36), and the evolutionary investment by the GHVs to subvert STAT3 signaling, we sought to further dissect the role of STAT3 in GHV latency establishment *in vivo*.

We previously used a mouse model in which STAT3 is ablated specifically in the CD19+ B cell compartment to discover that STAT3 is a crucial host determinant of MHV68 latency establishment in B cells *in vivo* (40). In the present study, we more closely examined virus latency in GC subsets and virus-specific adaptive immune responses in two independent strains of *CD19^cre/+^Stat3^f/f^* mice to eliminate possible strain variation. We identified disordered GC architecture in B cell-specific STAT3 knockout mice, along with reduced virus-specific antibody production, concomitant with a heightened CD8+ T cell response. This aberrant immune response in the absence of STAT3 signaling in B cells led us to use a mixed bone marrow chimera approach wherein mice are reconstituted with hematopoietic cells from WT and B cell-specific STAT3 KO mice. Infection of mixed bone marrow chimeric mice revealed that B cells lacking STAT3 could not compete with their wild-type (WT) counterparts to support GHV latency. To gain mechanistic insight into the transcriptional consequences of STAT3 loss, we sorted STAT3 WT and KO GC B cells, with or without infection, and quantified gene expression profiles with RNA sequencing. We identified a pro-viral role for STAT3 in dampening the interferon response.

## RESULTS

### Loss of STAT3 in B cells disrupts gammaherpesvirus latency and leads to abnormal GC structure

Mice with *loxP*-flanked exons of *Stat3* (*Stat3^f/f^)* that are crossed with mice expressing Cre recombinase under the control of the endogenous *CD19* promoter exhibit loss of STAT3 only in CD19+ B cells (41). Our lab previously reported that the conditional knockout (KO) of STAT3 in B cells led to a severe defect in the establishment of MHV68 latency in the spleens of mice after intranasal or intraperitoneal infection (40). In this study, we compared MHV68 latency in two strains of B cell-specific STAT3 knockout (KO) mice that differ by the location of *loxP* sites that flank exons of the *Stat3* gene (Fig. 1A) (42, 43). Consistent with our previous report for the *CD19^cre/+^Stat3^f/f^*-1 mice, the CD19+ B cell population of the *CD19 ^cre/+^Stat3^f/f^*-2 mice lacked detectable levels of STAT3 protein by immunoblot (Fig. 1A). For brevity, *CD19^+/+^Stat3^f/f^*mice will be referred to as WT mice, and their littermate *CD19^cre/+^Stat3^f/f^* mice will be referred to as B cell-STAT3 KO mice. STAT3 tyrosine 705 (Y705) phosphorylation is the classical indicator of STAT3 activation, and this activation has been demonstrated in response to human GHV infection in cell culture with endothelial, dendritic, and B cell models (44-47). We used intracellular flow cytometry staining of splenocytes from infected mice and revealed an increase in STAT3-Y705 phosphorylation in B cells from the spleens of infected WT mice compared to the littermate B cell-STAT3 KO mice (Fig. 1B).

**Fig. 1.**
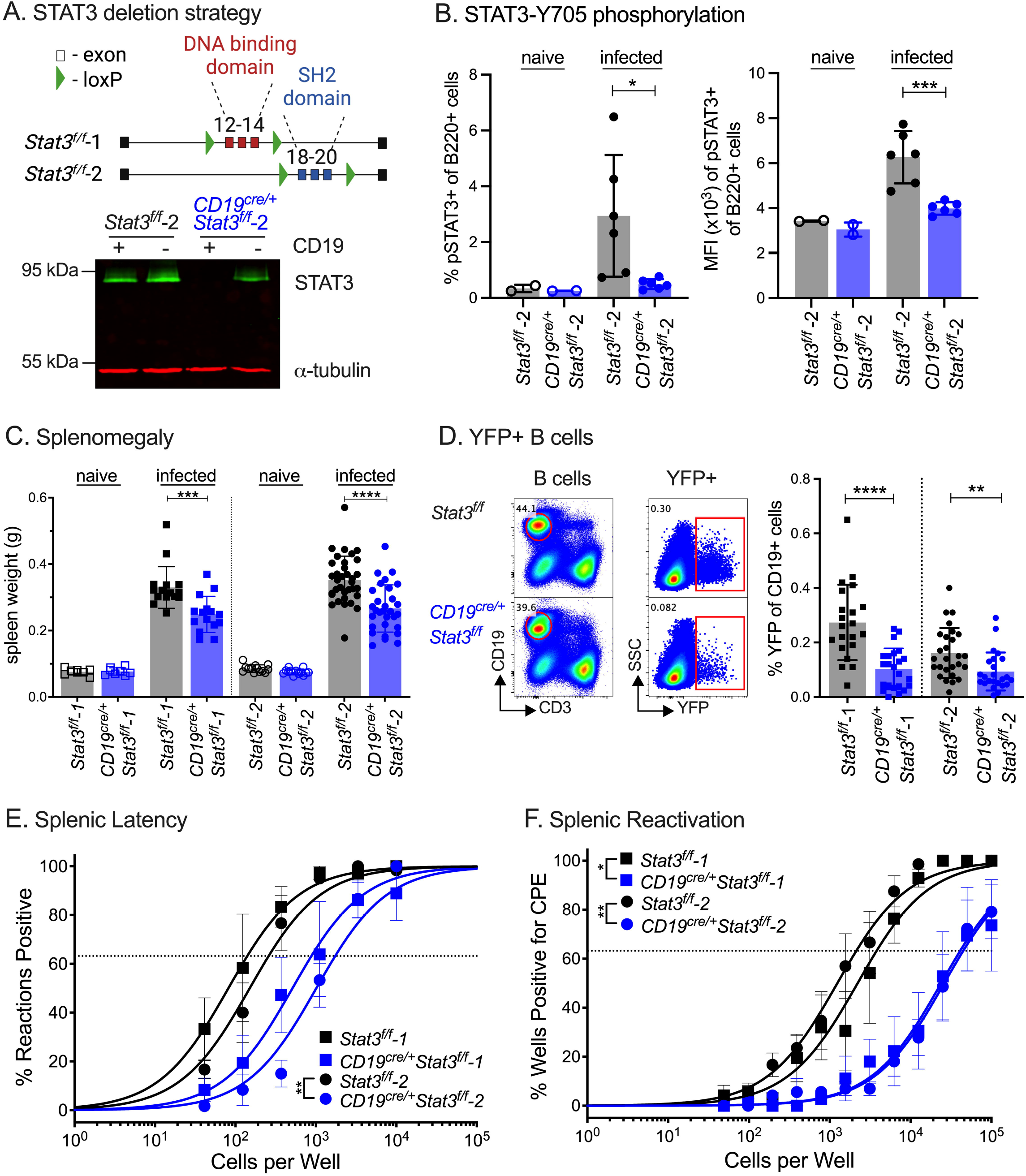
**STAT3 is necessary for the efficient establishment of latency in B cells of infected mice.** (A) Schematic of *loxP*-flanked exons of the *Stat3* locus in two strains of *CD19*^cre/+^*Stat3^f/f^* mice. Immunoblot confirmation of STAT3 loss in CD19+ B cells of *CD19*^cre/+^*Stat3^f/f^*-2 mice, but not CD19- non-B cells, as previously described for *CD19*^cre/+^*Stat3^f/f^* -1 mice (40). (B-F) *Stat3^f/f^* and *CD19^cre/+^Stat3^f/f^* mice were infected with 1,000 PFU MHV68-H2bYFP by intraperitoneal (i.p.) inoculation and evaluated at 16 days post-infection (dpi). (B) Intracellular staining of STAT3-Y705 phosphorylation in B220^+^ B cells of naive or infected mice evaluated as frequency of pSTAT3+ B220+ cells (left panel) and mean fluorescence intensity (MFI, right panel). (C) Weights of spleens from naive or infected mice. (D) Flow cytometry gating strategy (left panel) of infected (YFP+) B cells (CD19+CD3-) and enumeration of frequencies in each strain of mice (right panel). For B-D, each symbol represents an individual mouse; bars represent mean ± SD. Data shown represents two to three independent experiments performed with three to seven mice per infected group and one to two mice for naive groups. (E) Single-cell suspensions of spleen cells were serially diluted, and the frequencies of cells harboring an MHV68 genome were determined using a limiting dilution PCR analysis. (F) Reactivation frequencies were determined by *ex vivo* plating of serially diluted cells on an indicator monolayer. Cytopathic effect (CPE) was scored three weeks post-plating. For the limiting dilution analyses (E, F), curve fit lines were determined by nonlinear regression analysis; frequency values were determined by Poisson analysis, indicated by the dashed line. Symbols represent the mean ± SEM. Statistical significance was evaluated by two-tailed unpaired t test (B, C, D) or paired t test of the calculated frequencies (E, F). ∗p < 0.05, ∗∗p < 0.01, ∗∗∗p < 0.001, ∗∗∗∗p < 0.0001.

We previously reported that the loss of STAT3 in B cells did not affect acute replication of MHV68 in the lungs of mice after intranasal infection, but led to such a profound defect in latency establishment in the spleen that infected B cells were barely at the limit of detection (40). Here, we aimed to investigate the phenotype of infected B cells in the absence of STAT3, requiring us to use the more permissive intraperitoneal (IP) route of infection to enable sufficient infected cells for analysis. At 16 dpi, the peak of the early phase of splenic latency, infected mice exhibit an enlargement of the spleen that follows colonization of that tissue with MHV68 (4). Spleens from both strains of B cell-STAT3 KO mice were significantly reduced in mass upon infection, compared to the four-fold increase in mass observed upon infection of WT animals (Fig. 1C). The MHV68-H2bYFP recombinant virus utilizes a CMV immediate early promoter, driving constitutive expression of a histone 2b-YFP fusion protein and enabling direct analysis of infected cells (21). Consistent with the defect in splenomegaly, the frequency of B cells that expressed the YFP viral reporter gene determined by flow cytometry was reduced by three- and two-fold in the spleens of the strain 1 and strain 2 B cell-STAT3 KO mice, respectively, when compared to their WT littermates (Fig. 1D). This defect in latency was confirmed by a limiting dilution nested PCR assay to quantify the frequency of B cells harboring the viral genome. A 6.6-fold defect in the frequency of infected splenocytes that harbor the MHV68 genome was observed for *CD19^cre/+^Stat3^f/f^*-2 mice (1 positive event per 1,688 cells, 1/1,688) compared to *Stat3^f/f^*-2 mice (1/256) (Fig. 1E). Furthermore, in a limiting dilution reactivation assay, a ten and twenty-fold defect in spontaneous reactivation was observed upon explant of splenocytes from strain 1 and strain 2 of B cell-STAT3 KO mice compared to their WT counterparts, respectively (Fig. 1F). This reactivation defect largely mirrors the differences in latency. These results confirm the importance of STAT3 in B cells for the efficient establishment of MHV68 latency in two strains of *CD19^cre/+^Stat3^f/f^*mice.

The GC reaction represents an anatomical structure within secondary lymphoid organs in which activated B cells undergo clonal selection and affinity maturation. Additionally, most MHV68-infected cells express GC B cell markers at the peak of latency (21). Thus, we further examined GC B cell infection and latency in B cell-STAT3 KO animals. Compared to WT littermates, B cell-STAT3 KO exhibited increased frequencies of GC B cells (defined here as GL7+CD95hi of CD19+ cells) on 16 dpi compared to WT littermates (Fig. 2A). Although there was a reduction in the frequency of YFP+ B cells in B cell-STAT3 KO mice (Fig. 1D), we observed no significant difference in the frequency of YFP+ cells that displayed a GC phenotype (Fig. 2B). GC B cells can be roughly subdivided into rapidly proliferating centroblasts (CXCR4+CD86-) in the dark zone, and mature, non-dividing centrocytes (CXCR4-CD86+) in the light zone. By flow cytometry, infected WT mice had a 2:1 ratio of respective centroblast:centrocyte populations (Fig 2C). However, in the absence of B cell-STAT3, we observed increased frequencies of centrocytes in both total B cells and YFP+ GC B cell populations that resulted in a ratio closer to 1:1 (Fig. 2C).

**Fig. 2.**
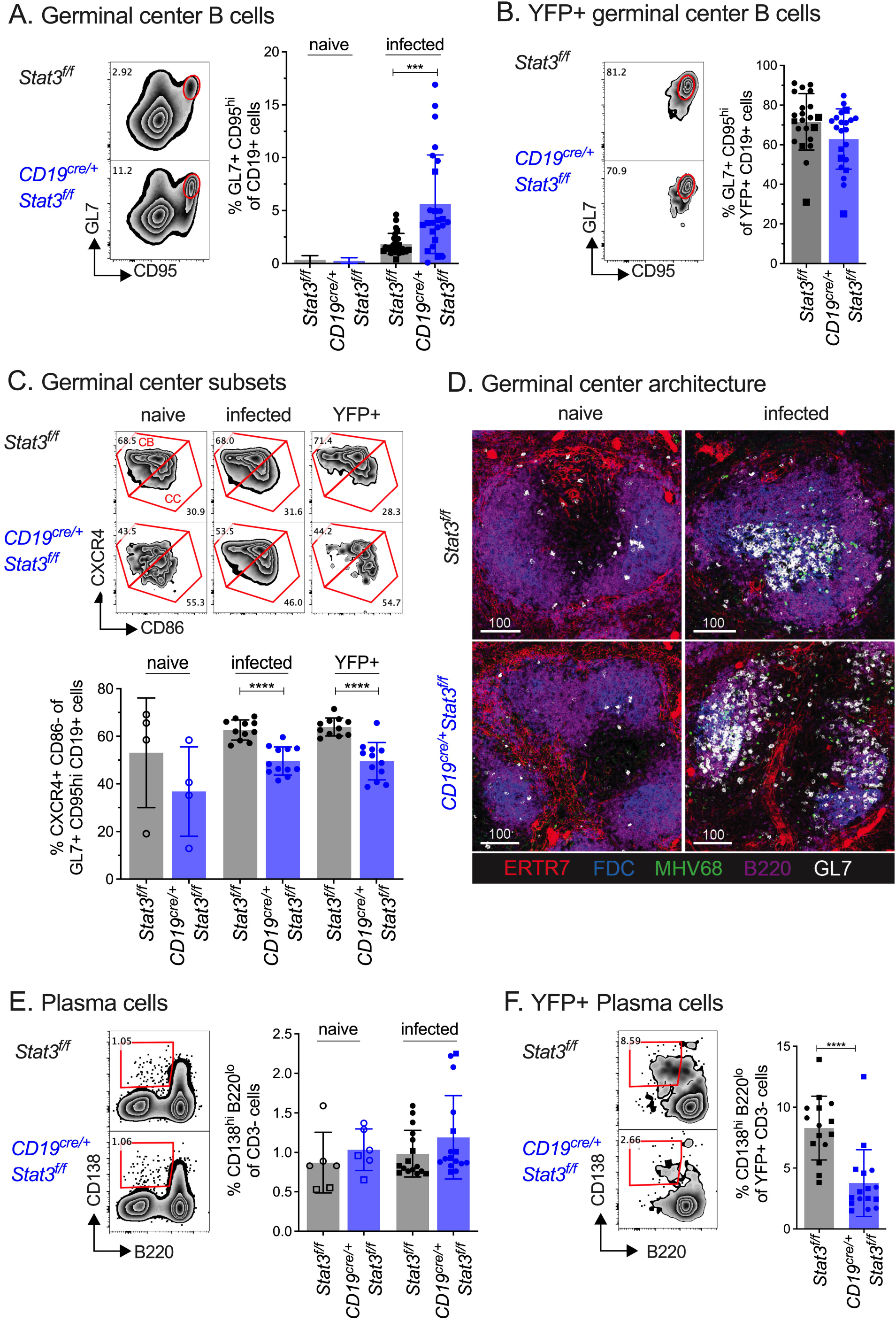
**STAT3 loss in B cells leads to an aberrant germinal center response upon infection**. *Stat3^f/f^* (WT) and *CD19^cre/+^Stat3^f/f^* (B cell-STAT3 KO) mice were infected with 1,000 PFU MHV68-H2bYFP by i.p. inoculation and evaluated at 16 dpi. (A) Flow cytometry gating strategy (left panel) and quantitation (right panel) of the frequencies of germinal center (GC) B cells (GL7+CD95+ of CD19+CD3-). (B) Flow cytometry gating strategy (left panel) and quantitation (right panel) of the frequencies of infected YFP+ cells bearing GC markers. (C) Flow cytometry gating strategy (upper panel) of dark zone centroblasts (CXCR4+CD86-) and light zone centrocytes (CXCR4-CD86+) as a frequency of GC B cells from naive or infected mice, and infected YFP+ B cells from infected mice. Quantitation of the frequencies of centroblasts of GC B cells from naive or infected mice, and infected YFP+ B cells (lower panel). (D) Confocal microscopy of frozen spleen sections from naive mice (left panels) or (right panels) from WT animals (*Stat3^f/f^*-2, top panels) or *CD19^cre/+^Stat3^f/f^*-2 mice (bottom panels) 16 dpi to identify stromal cells (ERTR7, red), FDCs (blue), B220+B cells (magenta) and GC B cells (GL7, white), MHV68-H2bYFP infection (green). Scalebars = microns. (E) Flow cytometry gating strategy (left panel) and quantitation (right panel) of the frequencies of plasma cells (CD138^hi^B220^lo^ gated on CD3-). (F) Flow cytometry gating strategy (left panel) and quantitation (right panel) of the frequencies of infected YFP+ cells with plasma cell markers. Data shown represents the mean ± SD of two to three independent experiments performed with four to seven mice per infected group and one to two mice for naive groups. Each symbol represents an individual mouse. Square symbols represent *CD19^cre/+^Stat3^f/f^-1* mice while circle symbols represent *CD19^cre/+^Stat3^f/f^-*2 mice. Statistical significance was evaluated by two-tailed unpaired t test (A-C, E-F). ∗∗∗p < 0.001, ∗∗∗∗p < 0.0001.

To further explore the increase in GC B cells of infected mice lacking B cell-STAT3, we examined GC architecture in the spleens by immunofluorescence (Fig. 2D). MHV68 infection of WT mice has previously been shown to result in the formation of visible, compact GCs within the splenic follicles (21). Interestingly, the B cell follicles appeared smaller and more diffuse in B cell-STAT3 KO mice than in WT spleens, even in uninfected animals. Upon infection, GC B cells did not remain in characteristic compact foci in B cell-STAT3 KO mice, instead appearing dispersed throughout the follicle and overlapping extensively with the follicular dendritic cells (FDC) network. In both WT and B cell-STAT3 KO mice, most of the YFP+ cells also expressed the GC marker GL7. These data indicate that STAT3-KO in B cells alone led to a significantly altered splenic architecture.

B cells typically exit the GC as class-switched, long-lived memory B cells or antibody-secreting plasma cells. STAT3 coordinates the upregulation of *Prdm1,* encoding BLIMP-1, a master regulator of B cell differentiation into plasma cells in response to IL-21 (48, 49). Plasma cells are a source of MHV68 reactivation in the spleen (50). Therefore, we examined plasma cell frequency and infection in the B cell-STAT3 KO mice (Fig. 2E). The frequency of B220^lo^CD138+ plasma cells between WT and B cell-STAT3 KO mice was similar (Fig. 2E), but we observed a marked reduction (> 50%) in YFP+ B cells with a plasma cell phenotype in the B cell-STAT3 KO mice (Fig. 2F). Together, the increase in GC cells, altered GC structure, and decrease in YFP+ plasma cells clearly indicate that general GC defects occur in the absence of B cell-STAT3. Collectively, this deeper analysis of two strains of B cell-STAT3 KO mice confirmed the requirement for STAT3 to promote B cell latency and revealed shifts in B cell subsets of the GC at 16 dpi. Because we observed similar latency and immune responses in both strains of mutant STAT3 mice, we proceeded with *CD19^cre/+^Stat3^f/f^-*strain 2 mice for the remainder of the study.

### Absence of STAT3 leads to dysregulated B and T cell responses 42 dpi

Due to observed immune perturbations in B cell-STAT3 KO mice at the early phase of chronic infection, we analyzed the B and T cell responses at a later 42 d timepoint. Mice produce virus-specific antibodies in response to MHV68 infection through a CD4+ T cell-dependent process (51, 52). The serum IgG levels were comparable between WT and B cell-STAT3 KO mice (Fig. 3A), but there was a significant decrease in virus-specific IgG in the serum of B cell-STAT3 KO mice (Fig. 3B). Accompanying this decrease, serum from B cell-STAT3 KO mice had a significantly reduced neutralization capacity by plaque reduction assay (Fig. 3C).

**Fig. 3.**
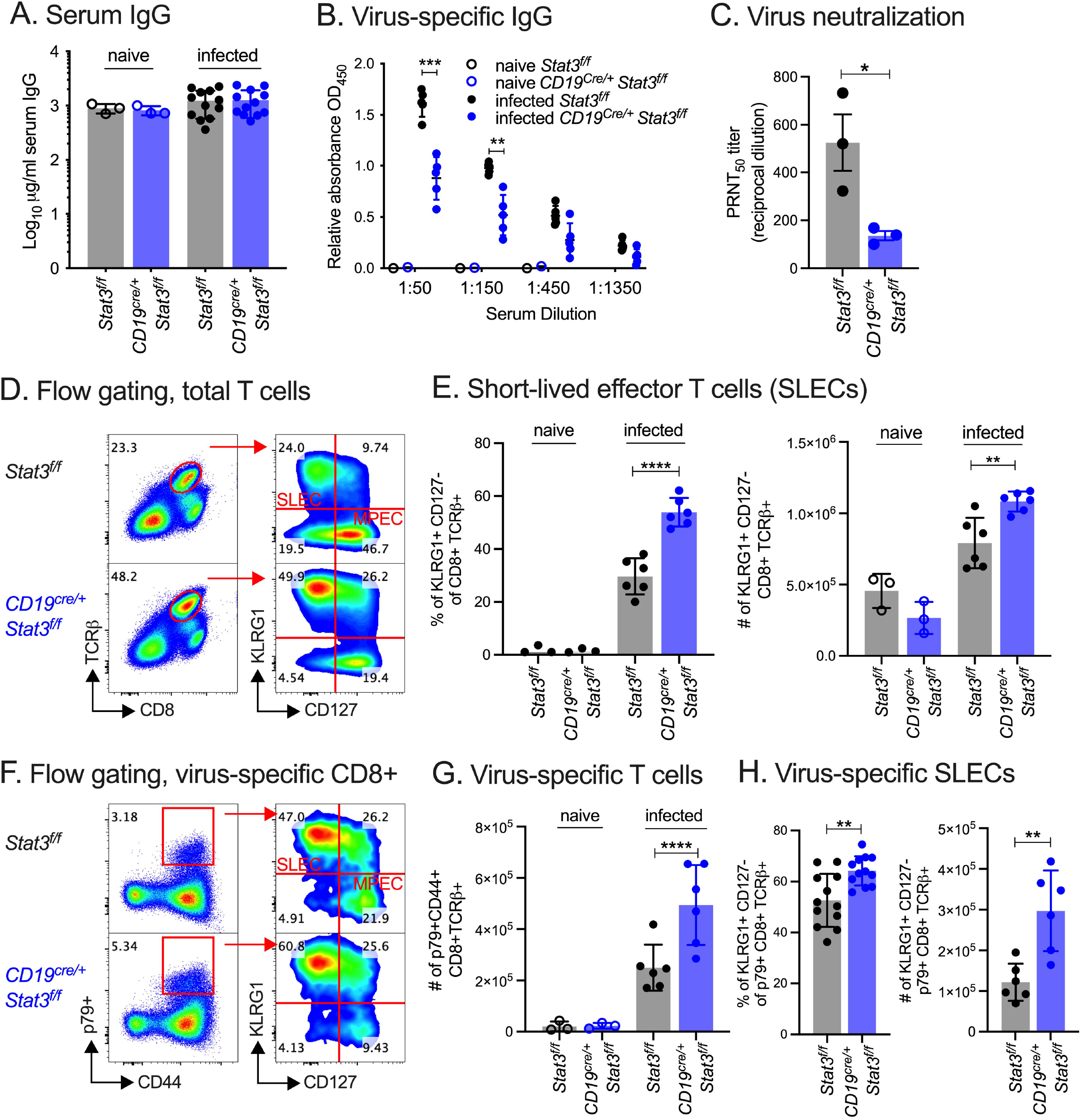
**Reduced antibody response and heightened T cell response to MHV68 infection in B cell-STAT3 knockout mice.** *Stat3^f/f^* (WT) and *CD19^cre/+^Stat3^f/f^*-2 (B cell-STAT3 KO) mice were infected with 1,000 PFU MHV68-H2bYFP by i.p. inoculation and evaluated at 42 dpi. (A-B) Total serum IgG (A) or virus-specific IgG (B) from naive or MHV68-infected *Stat3^f/f^* and *CD19^cre/+^ Stat3^f/f^* mice measured by ELISA. (C) Virus neutralization in serum as determined by a plaque reduction assay. The PRNT50 value is the dilution of serum to reach 50% neutralization of plaques. Symbols represent samples pooled from three independent experiments as biological replicates, each performed in technical triplicate. (D) Flow cytometry gating strategy for phenotyping short-lived effector CD8+ T cells (SLECs). (E) KLRG1+CD127- (SLEC) of CD8+TCRβ+ T cells by percentage of splenocytes (left panel) and total numbers per spleen (right panel). (F) Flow cytometry gating strategy for p79 tetramer+CD44+ (virus-specific) SLECs. (G) Total p79 tetramer+CD44+ (virus-specific) of CD8+TCRβ+ per spleen. (H) Analysis of virus-specific SLECs by percentage of spleen (left panel) and total numbers per spleen (right panel). Data shown represents the mean ± SD (A-B, E, G-H) of two to three independent experiments performed with four to seven mice per infected group and one to two mice for naive groups. For A-B, E, G-H, each symbol represents an individual mouse. Statistical significance was evaluated by two-tailed unpaired T test (B, C, E, G, H). ∗∗p < 0.01, ∗∗∗∗p < 0.0001.

We previously reported that the latency defect in B cell-STAT3 KO mice was maintained even during the maintenance phase of infection at 42 dpi (40), and we confirmed this for *CD19^cre/+^Stat3^f/f^*-2 mice (Supp. Fig 1A). T cells and effector cytokines such as interferon-γ control viral load during chronic infection with MHV68 (53). Activated, antigen-stimulated CD8+ T cells proliferate and differentiate into short-lived effector T cells (SLECs) or memory precursor effector T cells (MPECs) in response to infection. The B cell-STAT3 KO mice had an increase in SLECs by both percentage and total numbers when compared to their WT littermates 42 dpi (Fig. 3D-E). We quantified the number of CD44^hi^ CD8+ T cells binding to MHC class I tetramers presenting the immunodominant epitope p79 derived from MHV68 ORF61 (Fig. 3F). The spleens of B cell-STAT3 KO mice had twice the number of p79-specific CD8+ T cells (Fig. 3G), and a greater proportion of these virus-specific T cells were SLECs compared to WT mice (Fig. 3H). These unexpected aberrations in both the humoral and cell-mediated immune response confounded further mechanistic studies of B cell-STAT3 as a latency determinant and led us to develop an alternative mouse model to identify the B cell-intrinsic roles for STAT3.

### *In vivo* competitive infection model reveals B cell-intrinsic role for STAT3 in gammaherpesvirus latency establishment

Application of mixed BM chimeras to MHV68 infection has yielded important insight into the effects of B cell host determinants on infection (23, 25, 54). To better define the B cell-intrinsic role of STAT3, we generated mixed BM chimeras with STAT3 WT and KO B cells within the same animal. This creates an *de facto in vivo* competition model to analyze MHV68 latency in the context of the same adaptive immune response. To do this, CD45.2+ BM from *Stat3^f/f^tdTomato^stopf/f^*mice (WT STAT3 B cells) and from *CD19^cre/+^ Stat3^f/f^tdTomato^stopf/f^*mice (STAT3 KO B cells) was mixed 1:1 and transferred into CD45.1 recipients that had been irradiated for myeloblation (Fig. 4A). The *tdTomato^stopf/f^* reporter results in tdTomato red fluorescent protein production after Cre recombinase expression. In the *CD19^cre/+^Stat3^f/f^tdTomato^stopf/f^*mice, tdTomato expression marks B cell with STAT3 deletion, allowing for discrimination of STAT3 WT B cells and KO B cells. Sorted tdTomato+ B cells were analyzed by immunoblot to confirm the loss of STAT3 protein specifically in the CD19+tdTomato+ B cell population from *CD19^cre/+^Stat3^f/f^tdTomato^stopf/f^* mice, but not *Stat3^f/f^tdTomato^stopf/f^* WT B cell that lack Cre recombinase expression (Fig. S2A). TdTomato expression from CD19+ B cells and CD19 haploinsufficiency (*CD19^cre/+^*) did not impact MHV68 latency (Fig. S2B-F).

**Fig. 4.**
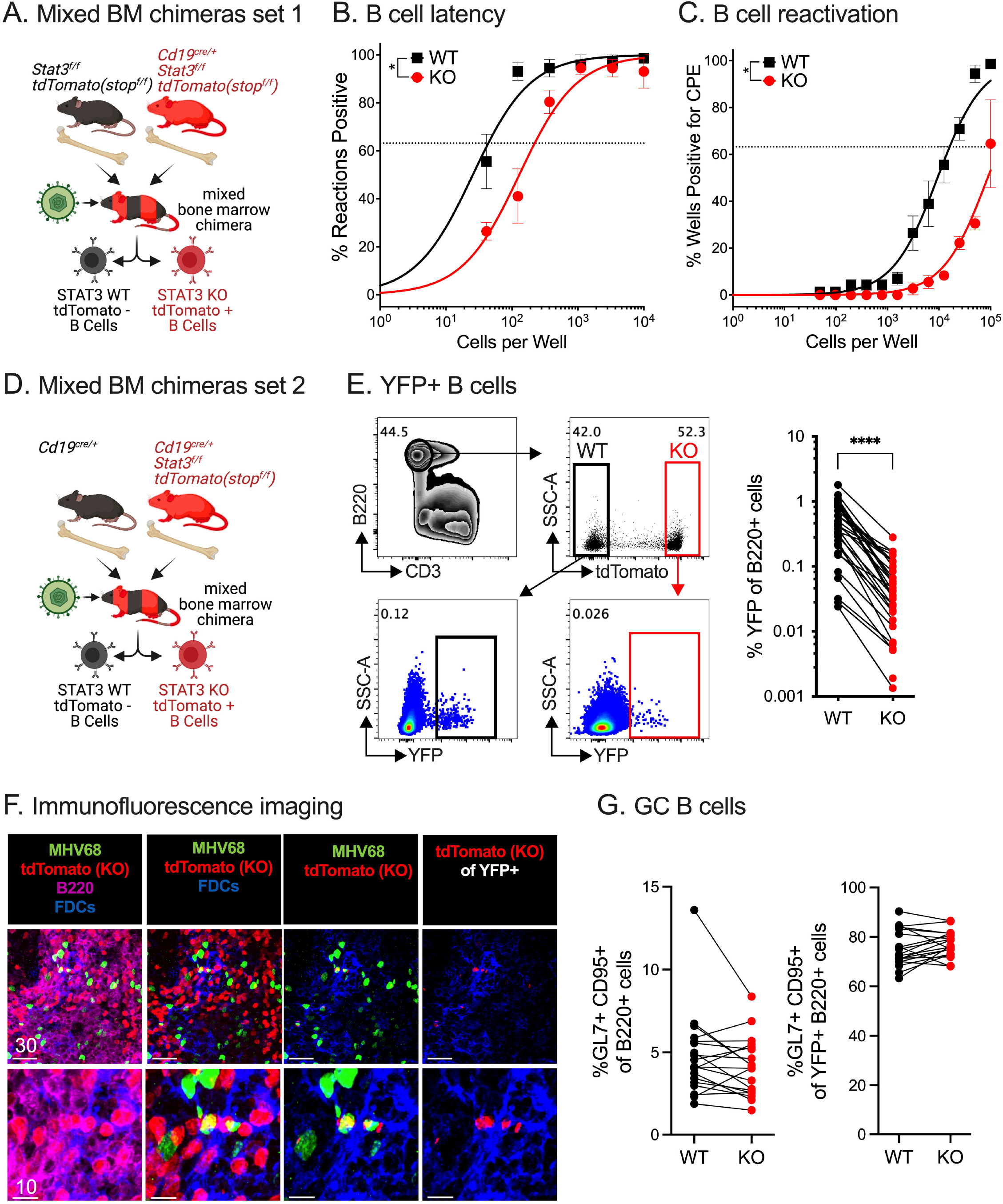
**Mixed bone marrow chimera models reveal intrinsic requirement for STAT3 in B cells for efficient establishment of latency with MHV68.** (A) Schematic depiction of the generation and infection of mixed bone marrow chimeras set 1 generated by reconstitution of CD45.1 C57/BL6 recipient mice with bone marrow from *CD19^+/+^Stat3^f/f^tdTomato^stopf/f^* and *CD19^cre/+^Stat3^f/f^tdTomato^stopf/f^* mice. (B-C) Mixed bone marrow chimeric set 1 mice were infected with 1,000 PFU MHV68-H2bYFP by i.p. inoculation and evaluated at 16 dpi. (B) Single-cell suspensions of sorted tdTomato- (WT) and tdTomato+ (STAT3 KO) B cells were serially diluted, and the frequencies of cells harboring MHV68 genomes were determined using limiting dilution PCR. (C) Reactivation frequencies were determined by *ex vivo* plating of serially diluted sorted tdTomato- (WT) and tdTomato+ (STAT3 KO) B cells on an indicator monolayer. CPE was scored three weeks post-plating. For the limiting dilution analyses (B, C), curve fit lines were determined by nonlinear regression analysis; frequency values were determined by Poisson analysis, indicated by the dashed line. Symbols represent the mean ± SEM. (D) Schematic depiction of the generation and infection of mixed bone marrow set 2 chimeras generated by reconstitution of CD45.1 C57/BL6 recipient mice reconstituted with bone marrow from *CD19^cre/+^*and *CD19^cre/+^Stat3^f/f^tdTomato^stopf/f^* mice. (E-G) Mixed bone marrow chimeric set 2 mice were infected with 1,000 PFU MHV68-H2bYFP by i.p. inoculation and evaluated at 16 dpi. (E) Flow cytometry gating strategy for infected (YFP+) STAT3 WT (tdTomato-) and STAT3 KO (tdTomato+) B cells (B220+CD3-) followed by quantitation of the frequency of infected B cells from mixed bone marrow chimeras. (F) Confocal microscopy of frozen spleen sections of spleens from chimeric mice harvested on day 16 post-infection. Staining = tdTomato (STAT3KO B cells, red); MHV68 (green); FDCs (blue); B220+ B cells (magenta). Top panels, lower magnification images; bottom panels higher magnification images of top panels. Far left panels, all colors; middle left omits B220 for easier visualization of tdTomato+ B cells; middle right panel, gated on the tdTomato cells infected with MHV68; far right panel omits MHV68 signal (green) for easier visualization of tdTomato+ B cells. Scalebars = microns. (G) Quantitation of STAT3 WT and STAT3 KO germinal center (GC) B cells of total B cells (left panel) or of YFP+ B cells (right panel). Data shown represents the mean ± SD of three independent experiments performed with six to seven mice per infected group. Each group of paired symbols represent the WT and KO B cell populations from one chimeric animal. Statistical significance was evaluated by paired t test; ∗p < 0.05, and ∗∗∗∗p < 0.0001.

At 16 dpi, splenocytes from the mixed BM chimeras were sorted to isolate tdTomato+ (STAT3 KO) and tdTomato- (STAT3 WT) B cells. Based on a limiting dilution, nested PCR for MHV68, KO B cells had a five-fold decrease in the frequency of genome-positive cells (average of 1/219) compared to their WT counterparts (average of 1/44) (Fig. 4B). The limiting dilution explant reactivation assay confirmed this decrease in latency establishment, but the frequency of spontaneous reactivation was not reduced further (Fig. 4C). These findings reinforce our initial findings (Fig. 1) and more rigorously demonstrate a B cell-intrinsic requirement for STAT3 in B cells for the efficient establishment of MHV68 latency.

To control for the haploinsufficiency of CD19 on the surface of *CD19^cre/+^* B cells, a second mixed BM chimera model was generated using donor *CD19^cre/+^Stat3^+/+^* BM in combination with BM from *CD19^cre/+^Stat3^f/f^tdTomato^stopf/f^*mice (Fig. 4D). Flow cytometric analysis of the chimeras revealed an eight-fold reduction in the frequency of MHV68-infected, YFP+ STAT3 KO B cells compared to their paired WT B cell counterparts (from 0.51% to 0.06%) in the same animal at 16 dpi (Fig. 4E). Confocal microscopy of frozen tissue sections of infected spleen sections from mixed BM chimeras identified STAT3 WT and KO (tdTomato+, red) B220+ B cells interspersed throughout the tissue (Fig. 4F). The ratio of STAT3 WT:KO B cells was consistent with the mean 39%:61% ratio of reconstitution for the WT and KO B cells, respectively, determined by flow cytometry. TdTomato+ KO B cells were rarely detected in the YFP+ population, in contrast to YFP+tdTomato- WT B cells. This is consistent with the failure of KO B cells to support latency in the splenic follicle based on limiting dilution PCR (Fig. 4B) and flow cytometric analysis of splenic single-cell suspensions (Fig. 4E). In contrast to the GC phenotype observed upon infection of mice that lacked STAT3 in all CD19+ cells (Fig. 2A), there was no difference in the frequency of GC B cells between the STAT3 WT and KO populations in the mixed BM chimera model (Fig. 4F-G, Fig. S3A-B). These findings substantiate STAT3 as a B cell-intrinsic latency determinant for GHV latency establishment in an *in vivo* competition model of infection.

STAT3 directly regulates host genes conducive to proliferation and plasma cell differentiation in response to cytokines in the GC microenvironment, but the STAT3-dependent transcriptional landscape of infected GC B cells has not been defined. Capitalizing on the ability to use flow cytometry to differentiate uninfected from YFP+ infected GC B cells, with and without STAT3 (tdTomato+), we sorted and collected specific subsets for RNA-sequencing. The CD45.2 donor cells were gated on tdTomato to sort the STAT3 KO (KO) from WT B220+ B cells, followed by non-GC (GL7-CD95-) and GC (GL7+CD95+) B cells (Fig. 5A). Last, the GC cells were sorted based on YFP to separate uninfected from infected GC cells. In a principal component (PC) analysis, the non-GC and GC samples demonstrated the most difference along PC1, followed by separation of YFP- and YFP+ samples (Fig. 5B). The infected YFP+ WT GC segregated from the YFP+ STAT3 KO GC samples in PC2. Hierarchical clustering of the top 350 variable genes across all samples revealed binodal clustering of genes based on their regulation in non-GC versus GC B cells (Fig. 5C).

**Fig. 5.**
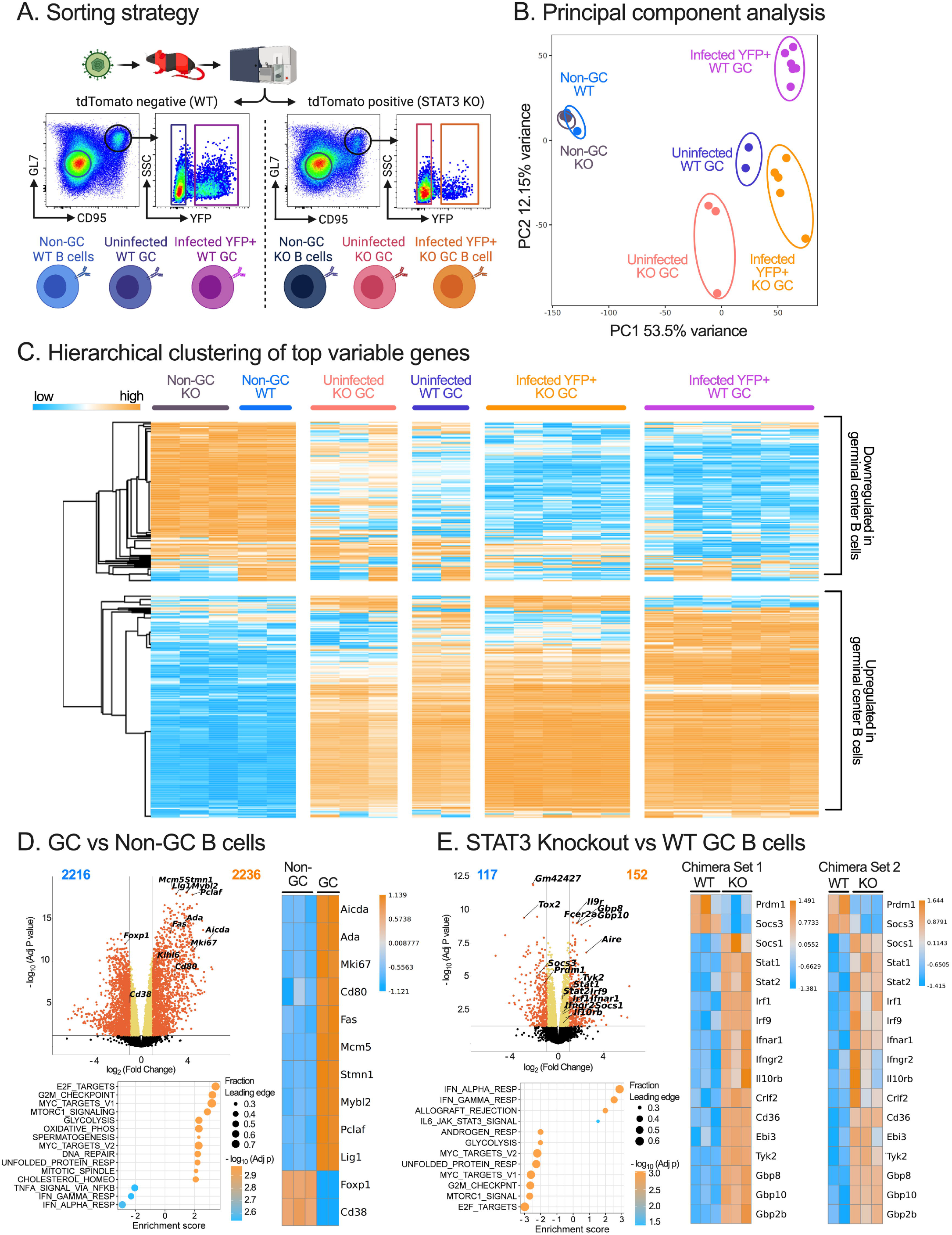
**RNA-sequencing of B cells from infected mixed bone marrow chimeric mice reveals transcriptional differences intrinsic to STAT3-ablated B cells.** (A) Schematic depiction of flow cytometry sorting strategy for B cell populations from mixed bone marrow chimeric mice. YFP+ infected and YFP- uninfected GL7+CD95+ germinal center (GC) cells were sorted and collected from tdTomato- and tdTomato+ populations. GL7-CD95- non-GC cells were also collected for tdTomato- and tdTomato+ populations. (B) Principal component analysis to analyze the clustering of RNAseq samples from the indicated flow sorted populations, mixed bone marrow chimeric set 1. (C) Hierarchical clustering of the top 350 variable genes across all samples, mixed bone marrow chimeric set 1. (D-E) For each, volcano plots with significance cutoffs displaying log_2_ fold change versus –log_10_ adj P value. Heatmaps display genes with adjusted P value <= 0.05 and at least 2-fold change between indicated comparisons. Gene Set Enrichment Analysis (GSEA) panels display HALLMARK gene sets with adjusted p value <= 0.05, |NES| >= 2. (D) Bioinformatic analysis comparing GC and non-GC B cells, mixed bone marrow chimeric set 1. Heatmap (right panel) of differentially regulated GC-specific genes curated from Broad Institute GSEA gene sets. (E) Bioinformatic analysis comparing uninfected STAT3 KO GC B cells to uninfected STAT3 WT GC B cells. Heatmaps (right panels) of DEGs in replicate RNAseq datasets from mixed bone marrow chimera set 1 and set 2; genes listed detail hallmark Interferonα Response and STAT3- regulated genes.

A comparison of WT GC to WT non-GC B cells in a volcano plot that highlighted genes with a minimum two-fold change and adjusted p value <0.05 identified the upregulation of GC- signature genes *Aicda, Mki67, Cd80,* and *Fas,* with downregulation of *Foxp1* and *Cd38* (55, 56). In a gene set enrichment analysis (GSEA), hallmark gene sets associated with proliferation such as E2F targets, G2M checkpoint, MYC targets, and MTOR signaling were positively enriched in the GC cells compared to the non-GC B cells, while interferon (IFN) α and IFNγ response pathways had a negative enrichment score (Fig. 5D). Next, KO GC cells were compared to their WT GC counterparts. Genes encoding B cell surface proteins IL-9R and CD23 (*Fcer2a*) were highly upregulated upon loss of STAT3, while known STAT3 target genes *Socs3* and *Prdm1* were downregulated (Fig. 5E, upper left panel). Hallmark gene sets for G2M checkpoint, E2F targets, and MYC proliferative pathways were negatively enriched in the KO GC cells in striking contrast to WT GC cells, highlighting the role of STAT3 as a driver of cell cycle and proliferation (Fig. 5E, lower left panel). Regarding upregulated gene sets, the IFNα and IFNγ response gene sets had the highest enrichment scores, indicating a heightened IFN profile in KO GC cells. An enhanced IFN response encompassed an increased expression of IFNα and IFNγ receptors and downstream signaling molecules *Stat1, Stat2, Irf9,* and IFN-stimulated genes (ISGs) including *Ebi3, Irf1, and GBP10*. The downregulation of proliferative pathways and heightened IFN responses in KO GC cells compared to WT GC cell counterparts was identified in both mixed BM chimera models (Fig. 5E, right panels). In summary, the loss of the master regulator STAT3 led to significant changes in the transcriptional landscape of GC cells, marked by the downregulation of genes driving proliferation and cell cycle progression and an upregulation of genes driving a type I and type II IFN response.

### Reduction of CD23 expression on infected B cells is influenced by STAT3

*Fcer2a* is the low-affinity IgE receptor commonly known as CD23, a surface marker used in combination with CD21 to delineate splenic B cell subsets. In the mixed BM chimera model, CD23 expression was notably increased in the KO B cells of infected mice (Fig. 6A). Higher CD23 expression in KO B cells manifested in a significant increase in the CD21^int^CD23^hi^ follicular subset (Fig. 6B, upper panel). WT B cells directly infected with MHV68 (YFP+) had much lower CD23 surface expression compared to KO cells (Fig. 6B, lower panel). The increase in follicular B cells based on heighted CD23 expression was also observed upon infection of the *CD19^cre/+^Stat3^f/f^* when compared to *Stat3^f/f^* mice (Fig. 6C-D). Taken together, MHV68 infection of B cells leads to a decrease in CD23 surface levels that are lost in the absence of STAT3.

**Fig. 6.**
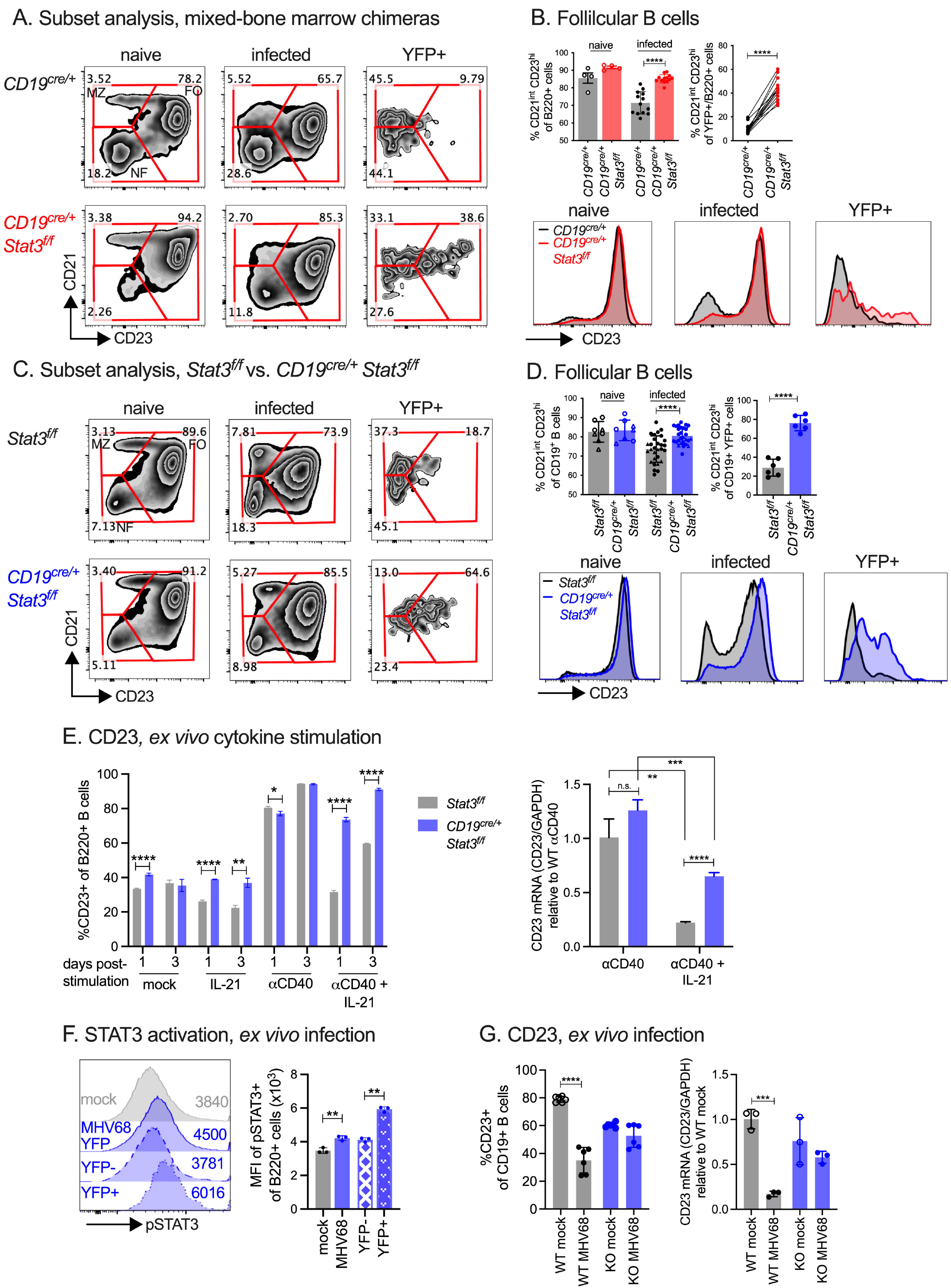
**MHV68-induced reduction of host CD23 surface expression lost in absence of STAT3.** (A, C) Flow cytometry gating strategy of splenic subset B cells for total B cells in naive or infected mice, or YFP+ B cells from infected mixed bone marrow chimeric set 2 mice (A) or *Stat3^f/f^* and *CD19^cre/+^ Stat3^f/f^* mice (C). (B, D) Quantitation (upper panels) of the frequency of follicular (CD21^int^CD23^hi^) B cells as gated in A and C, in total B cells from naive or infected mice (left upper panels, unpaired analysis) or YFP+ infected B cells from infected mice (right upper panels, paired and unpaired analyses). Histogram depiction (lower panels) of surface CD23 mean fluorescence intensity (MFI) of each population. (E) Frequency of B cells with surface CD23 expression (left panel) from *Stat3^f/f^* or *CD19^cre/+^Stat3^f/f^*mice upon *ex vivo* cytokine stimulation for indicated times. Fold-change of mCD23 transcript levels normalized to housekeeping GAPDH, relative to WT αCD40 treated samples (right panel). (F) Histogram depictions (left panel) and MFI (right panel) of intracellular staining for STAT3- Y705 phosphorylation in total B cells from mock or MHV68 H2b-YFP cultures, as well as YFP- or YFP+ infected B cells in the infected cultures. (G) Frequency of B cells with surface CD23 expression (left panel) from *Stat3^f/f^*and *CD19^cre/+^ Stat3^f/f^*B cells, mock infected or WT MHV68-infected, as indicated. Relative CD23 mRNA transcript levels (right panel) in mock and MHV68-infected B cell cultures. Fold-change of mCD23 transcript levels normalized to housekeeping GAPDH, relative to WT mock infected samples. Data shown represents the mean ± SD of two to three independent experiments performed with three to seven mice per infected group and one to two mice per naive groups (A-D) or one representative experiment of two experiments, each performed in triplicate (E-G). Statistical significance was evaluated by paired t test (B, F), two-tailed unpaired t test (D, G), and multiple unpaired t test (E). ∗p < 0.05, **p < 0.01, ***p < 0.001, ∗∗∗∗p < 0.0001.

IL-21 is a well-known activator of STAT3 that influences B cell activation and function in the GC (57, 58). IL-21 signaling in B cells is necessary for the efficient establishment of latency with MHV68 (23). Given the roles of CD40 and IL-21 signaling in the generation and maintenance of the GC reaction that supports MHV68 latency establishment, primary B cells were stimulated with CD40 and IL-21 alone, or in combination. Stimulation of primary WT B cells with IL-21 lead to STAT3-Y705 phosphorylation, (Fig. S4A), while activation of CD40 signaling led to the upregulation of CD23 in WT (*Stat3^f/f^*) B cells, as previously reported (59-61). At one and three days post-treatment, IL-21 in combination with CD40 reduced the surface levels of CD23 induced by CD40 alone on WT B cells by 36.7% (Fig. 6E, left panel). In contrast, this response was lost on KO (*CD19^cre/+^ Stat3^f/f^*) B cells), evidenced by only a 3.3% reduction in CD23 levels upon IL-21 in combination with CD40 stimulation compared to CD40 alone. At the transcript level, CD23 mRNA decreased with combination treatment of CD40 and IL-21 at 2 d post treatment in WT B cells as compared to CD40 alone (4.6 fold), but not to the same extent (1.9 fold) in KO B cells (Fig. 6E, right panel). This finding indicates that IL-21 suppresses CD23 expression in primary B cells at least in part through STAT3, consistent with CD23 transcript upregulation previously reported in an RNAseq profile of a diffuse large B cell lymphoma cell line treated with an shRNA targeting STAT3 (62).

MHV68 infection of primary B cells led to an increase in STAT3-Y705 phosphorylation in the YFP+ subset, indicating direct infection activates STAT3 (Fig. 6F). To discern if MHV68 infection leads to suppressed CD23 surface levels *ex vivo*, primary B cells from WT and B cell-STAT3 KO mice were infected and analyzed at 3 dpi. MHV68 infection led to a significant decrease in the percentage of CD23+ B cells in WT but not STAT3 KO B cell cultures (Fig. 6G). The downregulation of CD23 mRNA upon infection was lost in the absence of STAT3. These findings reveal a role for STAT3 in regulating CD23 in response to cytokines and GHV infection.

### Heightened interferon response in GC B cells lacking STAT3

MHV68 infection of WT GC cells led to a multitude of host transcriptional changes. Hierarchical clustering of differentially expressed genes comparing infected KO and WT GC cells revealed four distinct clusters of genes (Fig. 7A, Table S1). These genes clustered based largely on their infection status (clusters I and IV) and the presence of STAT3 (clusters II and III). Genes in cluster I such as *Il10* and *Sox5* were upregulated by the presence of the virus in WT GC, but to a lesser degree in infected STAT3 KO cells. Gene expression in clusters II and III were positively or negatively regulated by STAT3, respectively. Genes in cluster II include *Irf4, Prdm1, S1pr1,* and *Socs3*, while genes in cluster III include *Smad1* and *Eaf2.* Cluster IV genes such as *Fcer2a, Aire, Il9r,* and ISGs were downregulated by the virus in WT infection, but this virus-driven decrease was blocked in the absence of STAT3.

**Fig 7.**
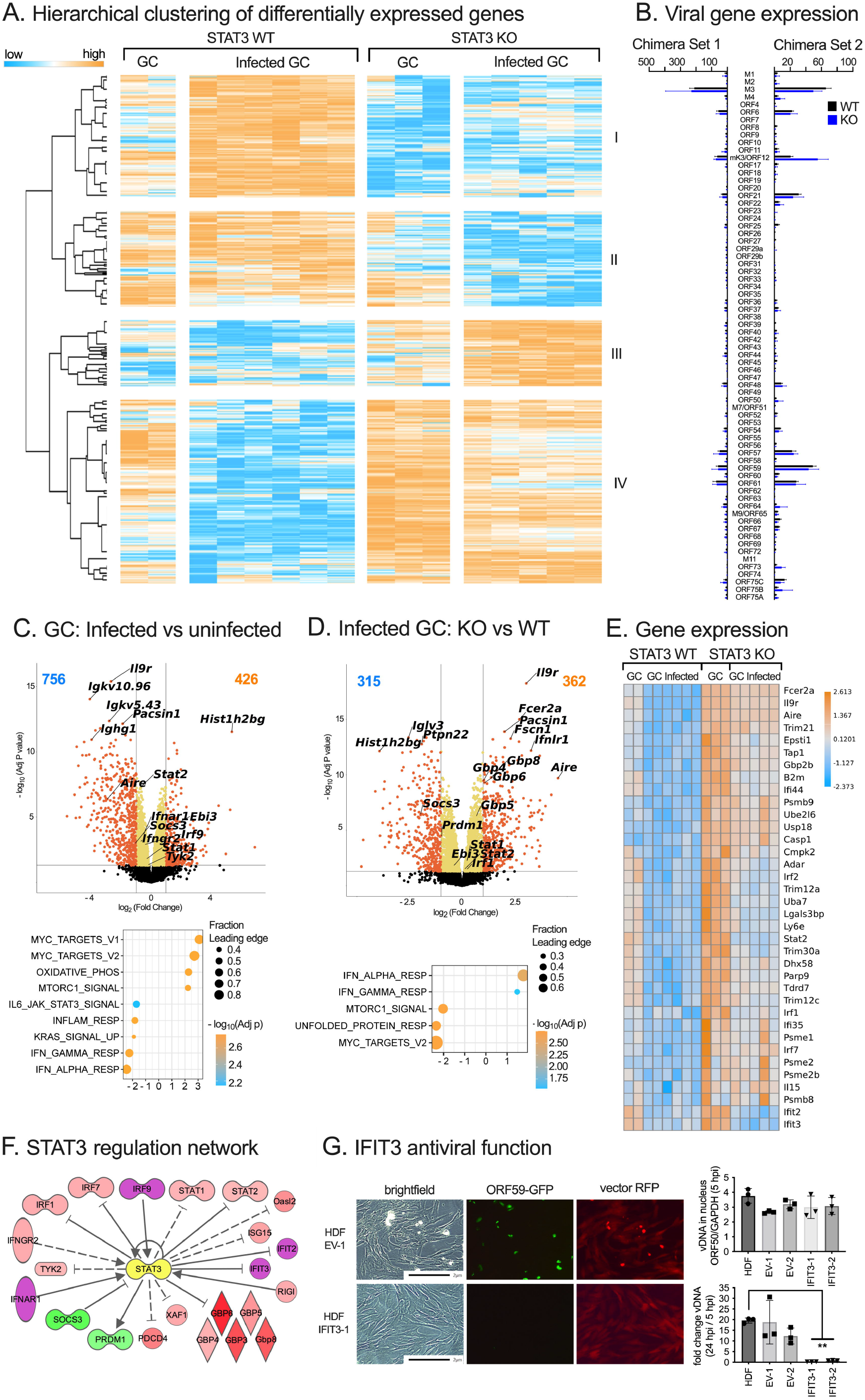
**Loss of STAT3 leads to a heightened interferon response in germinal center B cells.** (A) Hierarchical clustering of differentially expressed genes (DEGs) from the comparison of infected STAT3 KO and infected STAT3 WT GC B cells, separated into four distinct clusters of gene expression patterns. (B) Viral gene expression of infected YFP+ cells sorted from WT and KO GC B cells from the two mixed bone marrow chimera experiments. Bars represent the mean of the median-scaled CPM TMM values for each group of samples. (C-D) Volcano plots display log_2_ fold change versus –log_10_P value. GSEA panels display hallmark gene sets with adj p value <= 0.05, |NES| >= 2. (C) Bioinformatic analysis comparing infected WT germinal center (GC) and uninfected WT GC B cells. (D) Bioinformatic analysis comparing infected STAT3 KO GC B cells and infected STAT3 WT GC B cells. (E) Heatmap of gene expression for selected genes across uninfected and infected GC samples. The selected genes include leading edge genes from the Hallmark Interferonα Response gene set as revealed by our preranked GSEA of the infected KO versus WT GC B cell comparison, along with specific STAT3-regulated genes. These were filtered to retain only genes with adj p value <= 0.05 and at least 2-fold change comparing GC infected and uninfected WT and KO GC B cells. (F) STAT3 pathway analysis of DEGs identified in comparisons of infected KO and infected WT GC B generated by Ingenuity Pathway Analysis. Colors indicate relative expression changes; green, downregulation; red, upregulation, purple is a positive non-significant change. Symbols and lines indicate predicted role for STAT3; arrows, activation; straight lines, inhibition; solid lines; direct interaction; dashed lines, indirect interaction. (G) Telomerase (TERT)- immortalized human dermal fibroblasts (HDF) transduced with empty vector-RFP or IFIT3-vector-RFP, infected with an MHV68 ORF59-GFP reporter virus, MOI 3. Confocal microscopy for transduced cells (RFP) and early viral protein expression (GFP-tagged viral ORF59) at 24 hpi (left panel). Viral DNA isolated from the nucleus at 5 hpi (upper right panel) and fold change in viral DNA levels from 5 to 24 hpi by quantitative PCR (right panel) in parental HDF-TERT cells or HDF-TERT transduced with indicated constructs. Data shown represents the mean ± SD of one experiment performed with triplicate wells (G). Statistical significance was evaluated by one-way ANOVA (G). **p < 0.01.

Viral gene expression was tightly restricted in GC B cells, with very low levels of expression across the viral genome (Table S2). Although viral gene expression was minimal in GC cells at 16 dpi, there was a greater than four-fold higher expression level of genes with functions in immune modulation (*M3, mK3),* viral gene transactivation *(ORF57)* and viral DNA replication *(ORF6, ORF21, ORF59* and *ORF61*) compared to the mean expression level of all viral ORFs (Fig. 7B and Fig. S5). Notably, ORF6 and ORF61 are the source of immunodominant CD8 T cell epitopes in infected C57BL/6 mice (63). However, there were no significant changes in the viral gene expression profile of infected GC B cells in the absence of STAT3.

We and others previously reported a bias in the B cell Ig repertoire in the context of MHV68 infection; infected B cells exhibited immunoglobulin (Ig) heavy chain V gene usage that was non-overlapping with uninfected cells and displayed bias in lambda light chain (64, 65). Here, comparison of infected with uninfected WT GC cells revealed the downregulation of multiple Ig heavy chain and light chain V genes (Fig. 7C, upper panel). The predominant IgH V genes were distinct in the infected GC compared to their uninfected WT counterparts from both sets of BM chimera models (Fig. S6). Additionally, we confirmed that infected cells utilize lambda light chain more frequently.

Gene set enrichment analysis revealed positive enrichment for MYC targets and MTOR signaling, highlighting the increased expression of genes involved in cell cycle progression, such as *Psat1, Odc1, Bcat1, Pcna*, and *Mcm5.* Concurrently, there was a negative enrichment for genes involved in response to IFNα and IFNγ. This includes the downregulation of signaling molecules *Stat1, Stat2,* and *Irf9*, and ISGs *Dhx58, Ifit2, and Ifit3* (Fig. 7C, lower panel). Consistent with the known impacts of MHV68 in blocking host IFN responses (66-71), these findings reveal that MHV68 infection reprograms GC B cells *in vivo* to upregulate networks that promote proliferation while impairing antiviral IFN responses.

A comparison of infected KO and WT GC cells uncovered significant upregulation of *Fcer2a, Il9r, and Aire* in the absence of STAT3 (Fig. 7D, upper panel). As opposed to the upregulation of proliferative networks upon the infection of WT GC cells, MTOR and MYC gene sets were negatively enriched when comparing infected KO and WT GC cells, noted by the downregulation of cell cycle genes *Psat1, Odc1,* and *Bcat1* (Fig. 7D, lower panel). Additionally, genes of the IFNα and γ response pathways were upregulated in infected GC cells that lacked STAT3 (Fig. S7A-B), including the IFN signaling molecules *Stat1, Stat2,* and *Irf9,* and ISGs such as *Usp18* and *Gbp10* that were further validated by RT-qPCR (Fig. S7C). The heatmap of leading-edge genes for the IFNα response GSEA comparison of infected KO and WT GC cells in both mixed BM chimera datasets indicate that loss of STAT3 leads to a striking increase in expression of IFN signaling molecules and downstream ISGs (Fig. 7E).

Network analysis highlights multiple genes that were dysregulated in B cells lacking STAT3 (Fig. 7F). This includes known STAT3-responsive genes *Prdm1* and the suppressor of cytokine signaling factor 3, *Socs3.* STAT3 negatively regulates the type I IFN response by repressing the transcription of *Irf9*, as well as competing for binding partners with STAT1 (62, 72, 73). The loss of STAT3 in GC cells led to the upregulation of *Stat1*, *Stat2,* and *Irf9,* signaling molecules that drive type I and II IFN responses. ISGs induced in the absence of STAT3 include *Usp18, Oasl2*, multiple GBPs, and IFITs. IFIT1, 2, and 3 comprise a family of ISGs that form a complex and exhibit antiviral activity by binding viral mRNAs, inhibiting their translation (74, 75). IFIT3 overexpression blocks adenovirus replication through a MAVS/STING-mediated induction of the type I IFN response (76), but its role in herpesvirus infection has not been reported. Human dermal fibroblasts (HDF) were permissive for MHV68 infection based on the detection of the early lytic ORF59 replication factor at 24 hpi. In contrast, MHV68 ORF59 was not detected in HDF expressing IFIT3 (Fig. 7G). In addition, while equivalent levels of viral DNA were delivered to the nucleus at 5 hpi, an increase in viral DNA levels was blocked between 5 and 24 hpi in IFIT3-expressing HDF (Fig. 7G), consistent with a potent restriction of MHV68 at an early phase post-entry. The transcriptional profile indicates that uninfected GC cells lacking STAT3 are armed with numerous ISGs previously reported to restrict herpesviruses (77), in addition to IFIT3 as reported here. While virus infection dampens the interferon response in WT GC cells, it fails to do so in the absence of STAT3. These findings reveal pro-viral functions for STAT3 as a suppressor of both type I and II IFN responses and downstream ISGs that likely block early stages of viral infection.

## DISCUSSION

Here, we aimed to identify transcriptional and functional roles for STAT3 in B cells newly infected with a gammaherpesvirus *in vivo*. First, we verified a significant defect in latency establishment at 16 dpi in two strains of B cell-STAT3 knockout (KO) mice. This decrease in virus latency was accompanied by an increase in GC B cells and a reduction of infected plasma cells. Our analyses identified altered GC architecture in B cell-STAT3 KO mice, along with reduced virus-specific antibody production, and a heightened CD8 T cell response. This perturbed immune response led us to develop a mixed bone marrow chimera model wherein STAT3 KO B cells and WT B cells were both present as potential reservoirs of latency in the same host microenvironment. B cells lacking STAT3 were not able to compete with their wild-type counterparts to support GHV latency at 16 dpi. To dissect the transcriptional consequences of STAT3 loss, RNA sequencing was performed on sorted STAT3 WT and KO GC B cells, without and with MHV68 infection. Our analyses revealed multiple roles for STAT3, including a pro-viral role in dampening the interferon response.

We previously reported a requirement of STAT3 to support the efficient establishment of latency in a B cell-STAT3 KO mice model (40). Briefly, infection of B cell-STAT3 KO mice led to a reduction in splenic latency and in increase in GC B cells 16 dpi. Here, we expand upon these results in a second strain of STAT3 KO mice and report new findings of alterations in the virus-specific immune response. Immunization of STAT3-KO mice with OVA led to a notable decrease in GC B cells, short-lived GC structures, and a decrease in plasma cells (78). MHV68 infection resulted in an increase in total GC B cells in B cell-STAT3 KO mice, but these GCs were smaller and less dense compared to those in WT mice (Fig. 2D). MHV68 infection of mice leads to an increase in highly proliferative centroblasts compared to centrocytes within the infected GC population, as well as the total GC population (21, 23). In the absence of STAT3, we observed a reduced frequency of centroblasts in naive mice, as well as those infected with MHV68, with the ratio further skewed towards centrocytes in the infected GC population (Fig. 2C).

In the GC reaction, B cells differentiate and exit the GC as class-switched, long-lived memory B cells or antibody-secreting plasma cells. STAT3 coordinates the upregulation of *Prdm1*, encoding BLIMP-1, which is a master regulator of B cell differentiation to plasma cells in response to IL-21 (48, 49). These plasma cells are a source of MHV68 reactivation in the spleen (50) and long-lived plasma cells are a long-term latency reservoir in the bone marrow, which led us to examine plasma cells in the B cell-STAT3 KO mice. There was no difference in plasma cell frequency at 16 dpi, although there was a decrease in infected (YFP+) plasma cells in B cell-STAT3 KO mice. This phenotype was also reported for MHV68 infection of IL21R KO mice, where IL21 signaling is necessary for latency establishment (23). Regarding chronic infection, there was a significant reduction in virus-specific IgG (Fig. 3B) and virus neutralizing activity in serum (Fig. 3C) from B cell-STAT3 KO mice 42 dpi. The aberrant GCs and the reduction in centroblasts at 16 dpi along with the reduction in virus-specific antibodies suggests that GC cycling between dark and light zones may be disrupted in GCs of mice lacking STAT3 in B cells. These findings uncover a role for B cell STAT3 in GC processes and highlight effects likely downstream of the altered GC architecture and inappropriate B cell cycling in response to murine GHV infection.

The latency-associated M2 protein of MHV68 is critical for viral latency establishment in mice (79, 80). M2 induces high levels of IL-10 secretion in B cells and drives B cells to differentiate into plasmablasts *in vivo* (36-38). IL-10 is an anti-inflammatory cytokine that suppresses natural killer and T cell effector responses (81). An increase in virus-specific, short-lived effector CD8+ T cells (SLECs) was noted in the B cell-STAT3 KO mice at 42 dpi (Fig. 3H). The heightened virus-specific CD8+ T cell response may be a byproduct of the latency defect in B cell-STAT3 KO mice, driven by a reduction in MHV68 M2-driven production of the immunosuppressive cytokine IL-10. This finding is consistent with reports of increased virus-specific effector T cells in mice infected with a recombinant MHV68 virus that does not express the viral M2 protein (36).

Mixed BM chimeras have been employed to bypass immune defects caused by global deletion of signaling molecules CD40 and IL-21R or NF-κB p50 (23, 25, 54, 82). Here, BM from STAT3 B cell-competent and -ablated mice was mixed to create an *in vivo* model wherein STAT3 WT and KO B cells compete to support latency, in the context of the same microenvironment and immune response to infection. This model reveals that the loss of STAT3 leads to a marked reduction in MHV68+ B cells (Fig. 4B, E), confirming that STAT3 is required to promote B cell latency by GHV in the GC. This rigorous analysis supports STAT3 as a B cell-intrinsic host factor essential for latency establishment in primary B cells.

This is the first report of the transcriptional profile of MHV68-infected GC B cells. We capitalized on our ability to distinguish STAT3 KO B cells from WT counterparts based on Cre-inducible tdTomato expression in STAT3 KO B cells and the YFP reporter gene from MHV68- H2bYFP. FACS sorting for non-GC and GC subsets, with and without STAT3, with and without infection, followed by RNAseq enabled us to define the multitude of changes to the transcriptional landscape of GC B cells driven by the virus that are dependent on STAT3 (Fig. 5C). RNAseq of GC and non-GC B cells in the context of MHV68 infection revealed an enrichment for MYC targets by GSEA. MYC plays a major role in GC processes, regulating GC B cell entry and cycling between dark and light zones (83). Absence of MYC leads to reduced numbers of affinity-matured B cells within the GC. The impairment of MYC leads to the collapse of established GCs (84). Comparing STAT3 KO and WT GC B cells, there was a negative enrichment for MYC target genes by GSEA, consistent with MYC being a STAT3 target gene (62), and the role for STAT3 in MYC-driven proliferation (85, 86). Immunofluorescence of infected spleen sections revealed smaller follicles with less organized GCs in B cell-STAT3 KO mice (Fig. 2D). These findings call attention to the role for STAT3 in GC development after MHV68 infection.

The loss of STAT3 in GC B cells led to the downregulation of known STAT3 target genes *Prdm1* and *Socs3*, but also revealed the dysregulation of lesser studied genes such as *Il9r* and *Fcer2a* (Fig. 5E). IL-9 signaling promotes B cell exit from the GC and memory B cell differentiation (87). *Fcer2a*, also known as CD23, is the low-affinity IgE receptor. In humans, surface CD23 can be cleaved to a soluble form, where it can then bind membrane bound IgE and surface CD21 in complex to induce the production of secreted IgE (88). Along with its role in regulating IgE production, CD23 is upregulated by EBV and KSHV in lytic infection through the activity of viral transactivators EBNA2 and RTA respectively, and host RBP-Jκ (89-91). In the MHV68 latency model, CD23 was downregulated in total splenic B cells upon MHV68 infection in mice and upon *ex vivo* infection of primary B cell cultures, even more so in the directly infected cells (Fig. 6). CD40-ligation leads to the induction of CD23 on the surface of activated B cells (60, 61). We observed that IL-21 reversed CD40 induction only in the presence of STAT3, leading us to postulate that STAT3 is repressing CD23 expression through IL-21 signaling. *Fcer2a* was previously identified as a direct target that is negatively regulated by STAT3, in diffuse large B cell lymphoma cell lines (62). Future studies are needed to define the STAT3-dependent mechanism and consequence of MHV68-driven CD23 downregulation in B cells *in vivo*.

A consistent finding in the comparison of STAT3 WT and KO GC B cells was the positive enrichment for inflammatory and interferon (IFN) response pathways in the STAT3 KO cells. There was an increase in expression of genes involved in the IFNα and γ response pathways in the STAT3 KO B cells, both uninfected (Fig. 5E) and infected (Fig. 7D, E). Type I and II interferon responses restrict MHV68 latency through the induction of interferon stimulated genes (ISGs) which promote an antiviral state (4, 77, 92-94). Host factors that restrict MHV68 infection include those that promote IFN production like IRF3 and IRF7, and those that respond to IFN to directly induce ISG transcription like IRF1, STAT1, and STAT2 (66, 69, 95-98). Mice lacking STAT1, IFN-αβ receptor, or combination IFN-αβ and IFN-γ receptors succumb to infection with MHV68 (99), stressing the importance of the IFN response in combating the virus. MHV68 encodes numerous proteins to bypass the IFN response and promote replication and latency. MHV68 ORF11 and ORF36 interfere with IRF3 binding, blocking the transcription of IFNβ (67, 70). In addition, MHV68 ORF54 induces degradation of IFNAR1 to block IFN signaling (68), while the latency protein M2 impairs downstream IFN signaling by inducing the downregulation of STAT1 and STAT2 (66).

In the absence of STAT3, there was an upregulation of genes involved in type I and II IFN responses such as signaling molecules IRF9, STAT1, and STAT2 that comprise the interferon-stimulated factor 3 (ISGF3) transcriptional complex, as well as a multitude of ISGs such as *Oasl2, Usp18*, *Ifit3,* and the family of guanylate binding proteins (GBPs). Many of these ISGs have known antiviral function, such as human GBP1, encoded by *Gbp2b* in mice, which blocks KSHV capsid transport by disrupting cytoskeletal actin networks (100). An examination of the antiviral role of IFIT3 revealed an early block in MHV68 productive replication after viral genome delivery to the nucleus (Fig. 7G). STAT3 negatively regulates the type I and II IFN responses by directly repressing the transcription of IRF9, as well as competing for binding with STAT1, both leading to the reduction of ISGF3 transcriptional complex formation, and a reduction in IFN signaling (62, 72, 73). These results suggest that a heightened interferon response led to the reduction of splenic latency observed at day 16 in B cell-STAT3 KO mice (Fig. 1), as well as in STAT3 KO GC B cells in the mixed bone marrow chimera model (Fig. 4). This demonstrates STAT3 as a pro-viral host factor that dampens the antiviral IFN response in order to promote viral latency.

Our study highlights STAT3 signaling as a critical host pathway commandeered by GHVs to promote latency. Through sorting using reporter gene expression, we show that the requirement for STAT3 for MHV68 latency establishment is intrinsic to the infected B cell. STAT3 is activated in MHV68 infected cells, upregulating cell cycle genes to promote proliferation similar to EBV and KSHV (101, 102), while also subduing IFN responses (103) to circumvent clearance and promote latency. Understanding long-term latency of GHVs and how they utilize host factors and pathways to develop cancers is paramount. KSHV miRNAs block STAT3 and inhibition of STAT3 increases reactivation from latency (104). While there is a long-term latency defect in B cell-STAT3 KO mice 42 dpi, the role of STAT3 in the maintenance of MHV68 has yet to be uncovered. A recent study reveals a role for the STAT3-activating cytokine IL-16 in inhibiting reactivation through the STAT3-p21 axis (105). Though the use of JAK and direct STAT3 inhibitors show promise in clinical trials, deeper understanding of the role of STAT3 in viral persistence is required, as some GHV cancers may lose their dependency on STAT3 (106). This study provides a comprehensive examination into understanding signaling downstream of STAT3 in GHV infection *in vivo.* Future studies to delineate factors downstream of STAT3 in GHV infection and latency will reveal new avenues of treatment important for the prevention and treatment of GHV-driven malignancies.

## Acknowledgements

We thank the laboratories of Robert Yarchoan and Joseph M. Ziegelbauer for sharing critical equipment, as well as flow cytometry cores, animal facilities and support staff at SBU and the National Institutes of Health. We also thank Adrianus Van Der Velden, Pawan Kumar, Flaminia Talos and members of the Krug laboratory for helpful discussion.

## Author contributions

Conceptualization: C.H.H., N.C.R., L.T.K.

Methodology: C.H.H., N.C.R., L.T.K.

Software: C.H.H., T.J.M., L.T.K.

Validation: C.H.H., S.M.O., G.V.R., V.K., X.L., M.A.Z.

Formal analysis: C.H.H., S.M.O., X.L., M.A.Z., L.T.K.

Investigation: C.H.H., S.M.O., G.V.R., V.K., X.L., M.A.Z., Q.D., Y.Z., L.T.K.

Resources: A.C., C.K., B.S., K.M.M., P.H., H.D.H., J.C.F., L.T.K.

Data curation: C.H.H., T.J.M.

Writing – original draft preparation: C.H.H., L.T.K.

Writing – review and editing: C.H.H., L.T.K.

Visualization: C.H.H., S.M.W., V.K., X.L., M.A.Z., H.D.H., L.T.K.

Supervision: N.C.R., B.S., K.M.M., P.H., H.D.H., J.C.F., L.T.K.

Project administration: C.H.H., L.T.K.

Funding acquisition: N.C.R., H.D.H., L.T.K.

Declaration of interests: NONE

## METHODS

### Animal models

Strain 1 designates *Stat3^f/f^* mice with exons 12-14 flanked by *loxP* elements (42) and Strain 2 designates *Stat3^f/f^* mice with exons 18-20 flanked by *loxP* elements [B6.129S1- *Stat3^tm1Xyfu^*/J] (The Jackson Laboratory, Bar Harbor, ME) (43). *CD19^cre/cre^* mice [B6.129P2(C)- Cd19 ^tm1(cre)Cgn^/J] (Jackson) were crossed with Strain 1 and Strain 2 mice to generate *Stat3^f/f^CD19^cre/+^*-1 and -2 mice, respectively. *Stat3^f/f^* and *Stat3^f/f^CD19*^cre/+^ mice used in pathogenesis experiments were littermates derived from crossing *Stat3^f/f^* mice with *Stat3^f/f^* mice heterozygous for *CD19^cre^*; sexes were randomly assigned to experimental groups. *tdTomato^stopf/f^* mice [B6.Cg-*Gt(ROSA)26Sor^tm14(CAG-tdTomato)Hze^*/J] (Jackson) were crossed with *Stat3^f/f^ CD19^cre/+^* mice to generate *CD19^cre/+^Stat3^f/f^tdTomato^stopf/f^*mice. *CD19^cre/+^* mice were generated by crossing *CD19^cre/cre^* mice with C57BL/6 mice. Ly5.1 [B6.SJL-Ptprca Pepcb/BoyJ] (Jackson) were the recipient mice for bone marrow chimera studies. All mice were bred at the Stony Brook University Division of Laboratory Animal Research facility or the National Institutes of Health Division of Veterinary Resources, unless stated otherwise. Mice were housed in a specific pathogen–free environment, and all mouse experiments were performed in accordance with guidelines under protocols approved by the Institutional Animal Care and Use Committee of Stony Brook University and the NCI Animal Care and Use Committee.

### Mixed bone marrow chimera generation

Recipient Ly5.1 [B6.SJL-Ptprca Pepcb/BoyJ (Jackson) at six wks of age were gamma irradiated with 900 rads, in split dose. Next, mice were injected retro-orbitally with two million bone marrow cells isolated from donor *CD19^cre/+^Stat3^f/f^tdTomato^stopf/f^*and *Stat3^f/f^tdTomato^stopf/f^* mice at a ratio of 50:50 WT:KO bone marrow for the first set of chimeras, or with five million bone marrow cells isolated from donor *CD19^cre/+^Stat3^f/f^tdTomato^stopf/f^* and *CD19^cre/+^* mice at a ratio of 30:70 WT:KO bone marrow for the second set of chimeras. Mice were treated with acidified water two weeks prior to and after irradiation. Verification of chimerism was performed six weeks post transfer through collection of blood by submandibular bleed, followed by flow cytometry. The mean reconstitution ratio was 66% WT: 34% KO cells for the first set of chimeras and 39% WT: 61% KO for the second set of chimeras.

### Mouse infections and tissue processing

Recombinant MHV68 expressing H2bYFP fusion protein (rMHV68-H2bYFP) was used for the infection (107). Mice (8 to 12 weeks old) were infected by intraperitoneal injection with 1,000 PFU in 0.5 ml under isoflurane anesthesia. Mouse spleens were homogenized, treated to remove red blood cells, and passed through a 100-µm-pore-size nylon filter. For *ex vivo* experiments, flow sorting, and RNA-sequencing, splenocytes were subject to negative selection to enrich for B cells (Pan B cell isolation kit; STEMCELL, Vancouver, BC, Canada). Cells were counted using Vi-CELL BLU Cell Viability Analyzer (Beckman Coulter, Pasadena, CA). Blood was collected posthumously by cardiac puncture with gauge 27 needle. Serum was collected after a centrifugation at 20,000 x g for 20 mins at room temperature.

### Limiting dilution analysis of latency and reactivation

To determine the frequency of cells harboring the viral genome as an indicator of latency, single-cell suspensions were analyzed by single-copy nested PCR as previous described (108). To determine the frequency of cells harboring latent virus capable of reactivation upon explant, single-cell suspensions were plated in 12 serial 2-fold dilutions on a monolayer of MEF cells prepared from C57BL/6 mice and score for CPE three weeks after plating. Parallel samples were mechanically disrupted using a Mini-BeadBeater prior to plating on the monolayer of MEFs to release preformed virus and score for preformed infectious virus (108).

### Immunoblotting

Total protein lysate was harvested in lysis buffer (150 mM sodium chloride, 1.0% IGEPAL CA-630, 0.5% sodium deoxycholate, 0.1% sodium dodecyl sulfate, 50 mM Tris [pH 8.0]) supplemented with a protease inhibitor cocktail (Sigma, St. Louis, MO). Proteins were separated on 4-20% SDS-PAGE gels and transferred to a polyvinylidene fluoride membrane. Antibodies against STAT3 (clone K-15; Santa Cruz Biotechnology, Dallas, TX), and α-tubulin (clone B-5-1-2; Sigma) were detected using secondary anti-mouse (Rockland, Limerick, PA) or secondary anti-rabbit (Invitrogen, Grand Island, NY) antibodies by immunoblot analysis with an Odyssey Imager (Li-COR Biosciences, Lincoln, NE).

### Flow cytometry

For the analysis of B cells and T cells, 2 million splenocytes were resuspended in 50 µl of fluorescence-activated cell sorter (FACS) buffer (PBS with 2% fetal bovine serum) and blocked with TruStain fcX (clone 93; BioLegend, San Diego, CA). The cells were washed and stained to identify B cell subsets with fluorophore-conjugated antibodies against CD45.2 (clone 104), CD19 (clone 6D5), B220 (clone RA3-682), CD138 (clone 281-2), CD95 (clone 15A7), CD21 (clone 7E9), CD23 (clone B3B4), CXCR4 (clone L276F12), CD86 (clone GL-1), and CD3 (clone 17A2) or biotinylated antibody against GL7 (clone GL-7) that was detected using secondary streptavidin-conjugated allophycocyanin-cyanin7. T cell subsets were identified with antibodies against CD4 (clone GK1.5), CD8 (clone 53-6.7), CD44 (clone IM7), CD127 (clone A7R34), KLRG1 (clone 2F1/KLRG1), TCRβ (clone H57-597). H-2K(b)-p79 MHC-peptide complex was provided as biotinylated monomers by the NIH tetramer core facility and reconstituted with streptavidin-conjugated allophycocyanin. For STAT3-Y705 phosphorylation detection, cells were blocked with TruStain fcX before staining for surface markers. Following fixation and permeabilization with Fixation/Permeabilization kit (BD Biosciences, San Jose, CA), cells were permeabilized a second time in ice cold 100% methanol for 20 minutes. Cells were then stained with PE anti-STAT3 Phospho (Tyr705) antibody (Biolegend). For sorting, splenocytes were first enriched for B cells by depletion of non-B cells with magnetic microbeads (Pan B cell isolation kit; STEMCELL, Vancouver, BC, Canada) and then sorted using antibodies described above on a FACSAria III sorter (BD Biosciences) or MoFlo sorter (Beckman Coulter, Indianapolis, IN) into cold FACS buffer. All antibodies were purchased from BioLegend, Invitrogen, or BD Biosciences. The data were collected using a CytoFLEX flow cytometer (Beckman Coulter) and analyzed using FlowJo v10.8.1 (Treestar Inc., Ashland, OR). Dead cells were excluded based on Alexa Fluor™ 700 NHS Ester uptake. Doublets were excluded through forward scatter–height by forward scatter–area parameters.

### Enzyme-linked immunosorbent assay (ELISA)

To measure total IgG levels and antibody-specificity in the serum, plates were coated with either 2 µg/ml of donkey anti-mouse IgG (Affinipure; Jackson ImmunoResearch Laboratories, West Grove, PA), carbonate buffer (0.0875 M Na_2_CO_3_, 0.0125 M HCO_3_, pH 9.2) or 0.5% paraformaldehyde-fixed viral antigen–carbonate buffer and incubated overnight at 4°C. Coated plates were washed in PBS with 0.05% Tween 20 (PBS-T), and blocked in 3% milk– PBS-T prior to incubation with serial dilutions of serum or the mouse IgG standard (Invitrogen, Jackson ImmunoResearch) in 1% milk-PBS-T for 2 hrs at RT. IgG was detected by the use of horseradish peroxidase-conjugated donkey anti-mouse IgG (Jackson ImmunoResearch), SureBlue TMB (SeraCare, Milford, PA), and stop solution, and absorbance at 450 nm was read on FluoStar Omega (BMT Labtech, Cary, NC).

### Neutralization assay

Neutralization was tested by means of a plaque reduction neutralization test (PRNT) adapted from (52). Briefly, three-fold serum dilutions, starting from an initial concentration of 1:80 in Dulbecco’s modified Eagle medium (DMEM) supplemented with 10% fetal bovine serum (FBS), 100 U/ml penicillin, 100 μg/ml streptomycin, and 2 mM L-glutamine were incubated with 150 PFU of MHV68 on ice for one hr. The virus/serum mixture was then added to a sub-confluent BHK21 monolayer (4 x 10^4^ cells/well) plated the previous day in a 24-well plate, in triplicate. As a control, three wells received no-serum added virus. Plates were rocked every 15 mins for 1 h at 37°C. Infected cells were overlaid with 1.5% methyl cellulose solution in DMEM containing 2.5% FCS and incubated at 37°C for three to four days. Methylcellulose media was then aspirated, and cell monolayers were stained with a solution of crystal violet (0.1%) in formalin to facilitate identification and quantification of plaques. Percent neutralization was determined by comparison of the number of plaques in experimental wells compared to no-serum added control wells, and each data point was the average of three wells.

### Immunofluorescence

For confocal microscopy of frozen spleen sections, spleens were removed at the indicated time post-infection, fixed in periodate-lysine-paraformaldehyde fixative for 48 hr and then moved to 30% sucrose/PBS solution for 24 hr. Tissues were embedded in optimal-cutting-temperature (OCT) medium (Electron Microscopy Sciences) and frozen in dry-ice-cooled isopentane. 16-µm sections were cut on a Leica cryostat (Leica Microsystems, Buffalo Grove, IL). Sections were blocked with 5% goat, donkey, bovine, rat, or rabbit serum and then stained with a combination of the following Abs: ERTR7 (Abcam, Boston, MA), B220 (clone RA3-6B2, Biolegend), GL7 (clone GL-7, Thermo Fisher Scientific), CD21/35 (clone eBio4E3, eBioscience, San Diego, CA). Sections were incubated with secondary antibodies as needed and as controls, and images were acquired on a Leica SP8 or Stellaris Confocal microscope. To increase staining intensity of MHV68, spleens were stained with anti-GFP Ab (EPR14104, Abcam) and goat anti-chicken Alexa Fluor 488 (ThermoFisher Scientific, Waltham, MA). Images were processed and analyzed using Imaris software 8.0 (Oxford Instruments). Where indicated, the spots function of Imaris was used to identify and create spots for MHV68+ cells. Spots were masked on TdTomato expression inside the spot to reveal MHV68+ tdTomato+ cells, referred to as ‘gated’.

For immunofluorescence imaging of IFIT overexpressing HDFs, images were taken on an Olympus CLX41 microscope, with X-Cite 120 fluorescent system (Olympus, Center Valley, PA), with a mounted Progres Gryphax Kapella camera and Gryphax version 1.1.10.6 software (Jenoptik, Jena, Germany).

### Cell stimulation and ex vivo virus infection

For cytokine stimulations *ex vivo,* primary splenocytes were harvested from spleens of *Stat3^f/f^*, and *Stat3^f/f^CD19*^cre^ mice. Single-cell suspensions of splenocytes were subjected to negative selection to enrich for B cells (Pan B cell isolation kit; STEMCELL, Vancouver, BC, Canada). Primary B cells were plated in a 96 well plate at a concentration of 400,000 cells/ml, in 200 µl of primary B cell media (RPMI 1640 supplemented with 20% FBS, HEPES, non-essential amino acids, Sodium Pyruvate, Pennicillin/Streptomycin, L-Glutamine, β-mercaptoethanol). Cells were treated with anti-CD40 (BioLegend, 10 µg/ml) or murine IL-21 (25 ng/ml, PeproTech), alone or in combinations, as indicated. For *ex vivo* infection or primary B cells, cells were plated at a cell density of 10 million cells per ml in 200 µl in each well of a 24-well plate. rMHV68-H2bYFP was added at an MOI of 20, followed by centrifugation at 1500 x g for 1 hr at RT, and resuspended in 250 µl primary B cell media supplemented with 5 µg/ml LPS (Sigma).

For analysis of the impact of IFIT3 on infection, HDF-TERT cells (parental, empty vector, or IFIT3) (76) were infected with rMHV68-ORF59-GFP kindly provided by Craig Forrest, at an MOI 3 for two hrs, then washed twice with PBS, media was replaced, and cells were incubated until 5- and 24-hours post-infection (hpi). At 5 hpi, cell membranes were disrupted in 0.1%NP-40 in PBS, nuclei were spun out, and DNA was isolated. At 24 hpi, DNA was isolated from whole cells. qPCR for MHV68 immediate early gene ORF50 was performed on total DNA at 24 hrs and compared to nuclear DNA at 5 hpi.

### Quantitative reverse-transcription PCR

RNA was purified using RNeasy Plus Mini Kit (QIAGEN, Germantown, MD) following the manufacturer’s instructions. RNA was reverse transcribed to cDNA using the SuperScript IV First-Strand Synthesis system (Invitrogen) following the manufacturer’s instructions. Quantitative real-time PCR performed using PowerUp SYBR Green Master Mix (Applied Biosciences, College Station, TX) run on a QuantStudio 3 Real-Time PCR system (Applied Biosciences). Primers for GAPDH (F - CTCCCACTCTTCCACCTTCG, R - GCCTCTCTTGCTCAGTGTCC), and Fcer2a (F - CCAGGAGGATCTAAGGAACGC, R – TCGTCTTGGAGTCTGTTCAGG.

### RNA-sequencing

50,000 cells sorted on the markers of interest by flow cytometry were spun down, resuspended in 50 µl of TRIzol, and stored at -80C. GENEWIZ (South Plainfield, NJ) performed RNA extraction, quality control, library preparation, and Illumina sequencing.

The custom reference genome allowing quantification of both viral and host expression used in this alignment (“mm10_MHV68YFP_Krug”) consisted of the mouse reference (mm10/Apr. 2019/GRCm38) with a modified MHV68 sequence added as an additional pseudochromosome. This viral genome was prepared from the annotated herpesvirus genome (NCBI reference) with the addition of a CMV-driven Histone H2b-YFP fusion protein locus found in our mutant MHV68 virus strain, utilized to track individually infected cells. The custom gene annotations used for gene expression quantification consisted of a concatenation of the mm10 GENCODE annotation version M21 [99] and annotations of the MHV68 genome. The first annotation (“mm10_MHV68YFP_Krug_Ov”) measures expression using complete viral ORF annotations, including those regions that overlap. In the second annotation (“mm10_MHV68YFP_Krug_NoOv”), overlapping regions of the viral ORFs were removed to create a minimal, non-overlapping annotation. This second annotation was used to make conservative estimates of the expression of individual viral genes.

Raw sequencing files were aligned and counted using the CCR Collaborative Bioinformatics Resource (CCBR) in-house pipeline (https://github.com/CCBR/Pipeliner). Briefly, reads were trimmed of low-quality bases and adapter sequences were removed using Cutadapt v1.18 (http://gensoft.pasteur.fr/docs/cutadapt/1.18). Mapping of reads to custom reference hybrid genome described below was performed using STAR v2.7.0f in 2-pass mode (109, 110); Then, RSEM v1.3.0 was used to quantify gene-level expression (111), with quantile normalization and differential expression of genes analysis performed using limma-voom v3.38.3 (112). The data discussed in this publication, the custom genome reference FASTA, and both annotation GTFs have been deposited in NCBI’s Gene Expression Omnibus and are accessible upon request.

### Bioinformatics analysis and visualization

Data analysis and visualization were performed in the NIH Integrated Analysis Portal (NIDAP) using R programs developed on Foundry (Palantir Technologies) for normalization, differential expression analysis, gene set enrichment analysis (GSEA), and visualization. First, raw counts were imported and transformed to counts per million (CPM) and filtered to retain genes that had at least two samples with nonzero CPM counts in at least one group. Next, quantile normalization and batch correction across samples were used to account for factors that would prevent direct comparisons between samples. Heatmaps were created using the quantile-normalized, batch-corrected RNA-seq data. A principal component analysis demonstrating the within and between group variance in expression after dimensionality reduction was generated using NIDAP. Differential expression of genes (DEG) analysis utilized log_2_-counts per million (logCPM) transformed raw counts, implemented with the Limma Voom R package. Volcano plots were created from DEG analysis of each comparison highlighting significant fold changes (log_2_ Fold change) > 1 or < -1, with a significant p value (adj p value < 0.05). GSEA in NIDAP used the Limma Voom calculated t-statistic ranking to test comparisons against the Broad Institute GSEA hallmark curated gene sets.

Genes chosen for heatmap visualization represent differentially regulated genes that intersected with GC- (5D), STAT3- (5E), or viral infection- (5C) signature genes. Gene signatures were developed using the Molecular Signature Database (MSigDB, (113)) with GO annotations provided by the Gene Ontology resource. GO Databases: (114, 115). Germinal center signature developed using intersection of the following GO datasets: GO:0002314, GO:0002467, GO:0002636, and GO:0002634, accessed on 2022-12-15. Viral infection signature developed using intersection of the following GO datasets: Hallmark: MM3877, GO:0016032, GO:0039528, GO:0045071, GO:1903901, and GO:0002230. Hierarchical clustering heatmaps were generated using Morpheus, (https://software.broadinstitute.org/morpheus). Hierarchical clustering for all samples or 350 top variable genes were performed using a one minus pearson correlation metric, with complete linkage. CPM TMM values for viral ORFs were normalized using a scaling factor generated from median total viral counts of samples within a dataset. STAT3 network visualization was generated using curated data from QIAGEN IPA (QIAGENInc., https://digitalinsights.qiagen.com/IPA).

### Statistical analyses

Data were analyzed using GraphPad Prism software (Prism 8; GraphPad Software, Inc., La Jolla, CA). Significance was evaluated by unpaired two-tailed t test, paired t test, one-way ANOVA, or paired one-tailed t test of the log-transformed frequency values of samples from matched experiments, as noted. Using Poisson distribution analysis, the frequencies of latency establishment and reactivation from latency were determined by the intersection of nonlinear regression curves with the line at 63.2%.

## RESOURCE AVAILABILITY

### Lead contact and materials availability

Further information and requests for resources and reagents should be directed to and will be fulfilled by the lead contact, Laurie T. Krug.

### Data and code availability

Hybrid genome, raw sample reads, and BAM alignments are accessible upon request.

### Code

This paper does not report original code. Any additional information required to reanalyze the data reported in this paper is available from the lead contact upon request.

