## Supplemental Text for "B cell-intrinsic STAT3-mediated support of latency and interferon suppression during murine gammaherpesvirus 68 infection revealed through an *in vivo* competition model"

Supplementary Material

### Supplementary Methods

Mice

$tdTomato^{stopf/f}$ mice [B6.Cg-*Gt(ROSA)26Sor*$^{tm14(CAG\text{-}tdTomato)Hze}$/J] (Jackson) were crossed with $CD19^{cre/+}$ mice to generate $CD19^{cre/+f}tdTomato^{stopf/f}$ mice.

Immunoblotting

Antibodies against tyrosine 705-phosphorylated STAT3 (Cell Signaling Technology, Danvers, MA), and GAPDH (clone D4C6R, Cell Signaling Technology) were detected using secondary anti-mouse (Rockland, Limerick, PA) or secondary anti-rabbit (Invitrogen, Grand Island, NY) antibodies by immunoblot analysis with an Odyssey Imager (Li-COR Biosciences, Lincoln, NE).

Ex vivo cytokine stimulation

Cells were treated with murine IL-6 (25 ng/ml, PeproTech, Cranbury, NJ), murine IL-10 (25 ng/ml, PeproTech), or murine IL-21 (25 ng/ml, PeproTech), for 15 minutes prior to cell lysing to collect protein lysate.

Ig Repertoire Analysis

Ig repertoire was reconstructed from RNAseq datasets using TRUST4 algorithm (1) that annotated simultaneously heavy and light chain V genes on assembled contigs. All annotated reads were used to generate V gene heat maps.

Quantitative reverse-transcription PCR

Primers for Usp18 (F - CAGACGTGTTGCCTTAACTCC, R - ACTCCGAGGCACTGTTATCC), Gbp2b (F - ACCTGGAGACTTCACTGGCT, R - TTTATTCAGCTGGTCCTCCTGTATCC), Gbp8 (F

- CACACCCCACTAAACCAGAGC, R - TTCCATCAGGATTTGGTGAAGAC), Gbp10 (F - TTGTTGGATGGTCCCGTACT, R - GTGATTCTGTCCCGCCAG) used for RT-qPCR.

**Supplemental Figure Legends**

**SFig 1.**

*Stat3*$^{f/f}$ and *CD19*$^{cre/+}$*Stat3*$^{f/f}$-2 mice were infected with 1,000 PFU MHV68-H2bYFP by i.p. inoculation and evaluated at 42 dpi.

Single-cell suspensions of spleen cells were serially diluted, and the frequencies of cells harboring an MHV68 genome were determined using a limiting dilution PCR analysis. For the limiting dilution analyses, curve fit lines were determined by nonlinear regression analysis; frequency values were determined by Poisson analysis, indicated by the dashed line.

Data shown represents one experiment performed with four to five mice per infected group.

**SFig 2.**

(A) Immunoblot indicating the loss of STAT3 protein CD19+ sorted B cells from *CD19*$^{cre/+}$*Stat3*$^{f/f}$*tdTomato*$^{stopf/f}$. CD19+ populations from *CD19*$^{cre/+}$*Stat3*$^{f/f}$*tdTomato*$^{stopf/f}$ mice were sorted tdTomato+, indicating the expression of Cre recombinase.

(B-D) *tdTomato*$^{stopf/f}$ and *CD19*$^{cre/+}$*tdTomato*$^{stopf/f}$ mice were infected with 1,000 PFU MHV68-H2bYFP by intranasal inoculation and evaluated at 16 dpi. (B) Weights of spleens from naive or infected mice. (C) Single-cell suspensions of spleen cells were serially diluted, and the frequencies of cells harboring an MHV68 genome were determined using a limiting dilution PCR analysis. (D) Reactivation frequencies were determined by *ex vivo* plating of serially diluted cells on an indicator monolayer. Cytopathic effect (CPE) was scored at two- and three-weeks post-plating.

(E-F) *Stat3*$^{f/f}$*tdTomato*$^{stopf/f}$ and *CD19*$^{cre/+}$*Stat3*$^{f/f}$*tdTomato*$^{stopf/f}$ mice were infected with 1,000 PFU MHV68-H2bYFP by intranasal inoculation and evaluated at 16 dpi. (E) Single-cell suspensions of spleen cells were serially diluted, and the frequencies of cells harboring an MHV68 genome were determined using a limiting dilution PCR analysis. (F) Reactivation frequencies were determined by *ex vivo* plating of serially diluted cells on an indicator monolayer. Cytopathic effect (CPE) was scored at two- and three-weeks post-plating.

For the limiting dilution analyses (C, D, E, F), curve fit lines were determined by nonlinear regression analysis; frequency values were determined by Poisson analysis, indicated by the dashed line.

**SFig 3.**

Flow gating strategy for the frequency of total (A) or YFP+ (B) GC B cells (GL7+CD95+ of CD19+CD3-) from mixed bone marrow chimera set 2.

**SFig 4.**

Immunoblot confirmation of STAT3-Y705 phosphorylation in enriched CD19+ B cells with treatment of indicated cytokines. Samples were treated for 15 minutes before collection of total protein lysates.

**SFig 5.**

Viral gene expression from infected YFP+ cells sorted from WT and KO GC B cells from mixed bone marrow chimeras set 1 (A) or set 2 (B). Bars represent the mean of the median-scaled CPM TMM values for each group of samples. Data displayed on a log scale to showcase low viral gene expression.

**SFig 6.**

(A) Heat map showing utilization of top heavy and light chain V-genes in WT non-GC B cells, GC YFP- and GC YFP+ from mixed bone marrow chimera set 1 (A) and set 2 (B) RNAseq experiments. Only genes that are among top 15 V gene segments in each B cell subset are shown.

**SFig 7.**

Gene set enrichment analysis (GSEA) of the Hallmark Interferon $\alpha$ Response (A) or the Hallmark Interferon $\gamma$ Response (B). The enrichment plot for the comparison of infected KO and infected WT GC B cells is shown in the left panel, with the leading edge genes displayed as a heatmap for all GC samples in the right panel.

(C) RT-qPCR validation of select interferon stimulated genes in enriched B cells of naive and MHV68-infected *Stat3*$^{f/f}$ and *CD19*$^{cre/+}$*Stat3*$^{f/f}$-2 mice 16 dpi. Fold-change of indicated transcript levels normalized to housekeeping GAPDH, relative to the naive sample of each genotype.
