## Supplemental Figs1-7 for "B cell-intrinsic STAT3-mediated support of latency and interferon suppression during murine gammaherpesvirus 68 infection revealed through an *in vivo* competition model"

Supplementary Fig. 1

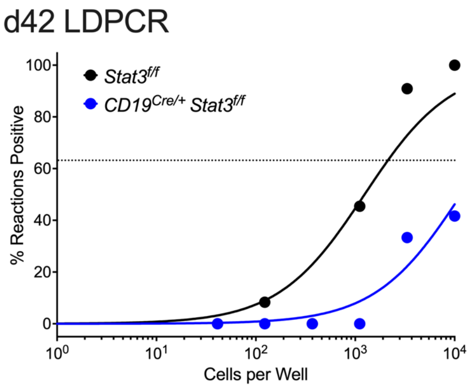

Supplementary Fig. 2

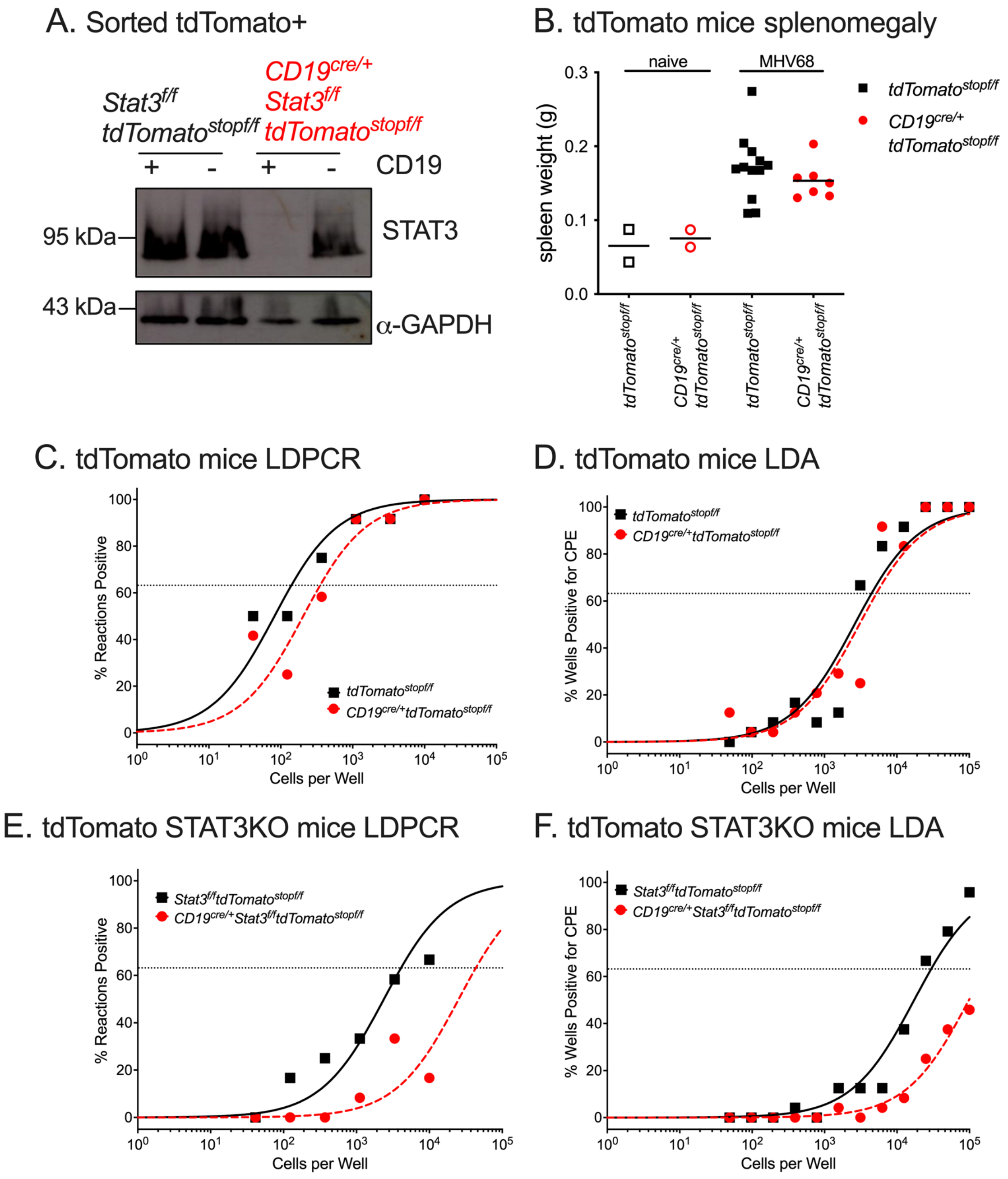

Supplementary Fig. 3

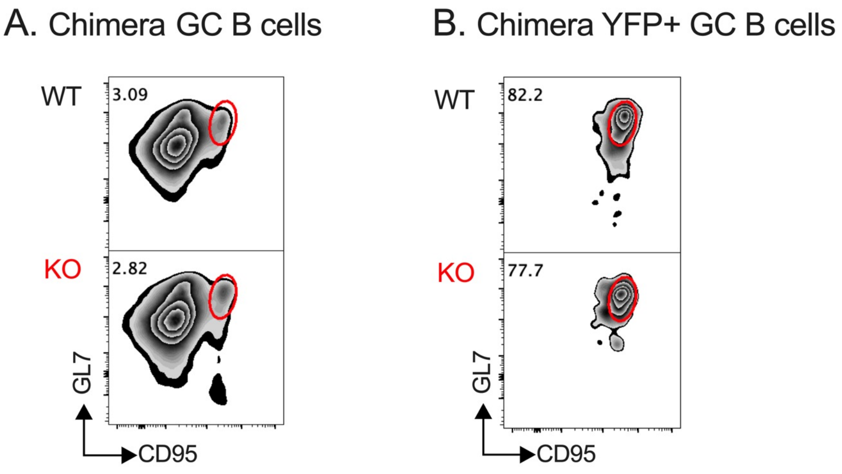

Supplementary Fig. 4

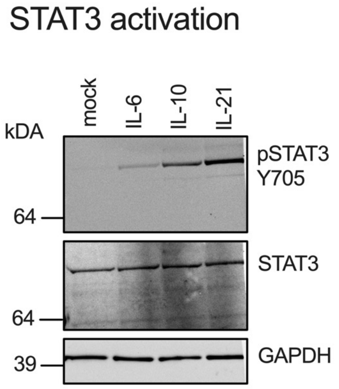

Supplementary Fig. 5

#### A. Viral gene expression (mBMC set 1)

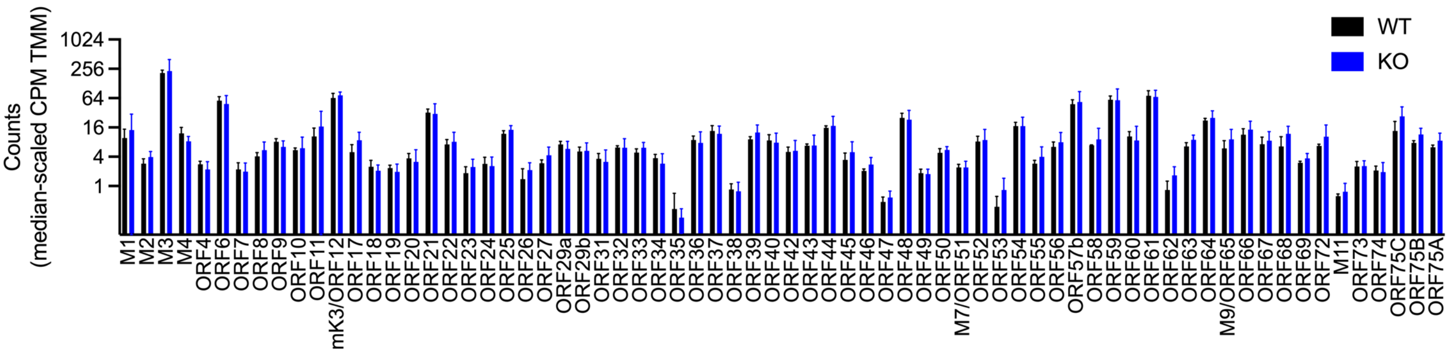


#### B. Viral gene expression (mBMC set 2)

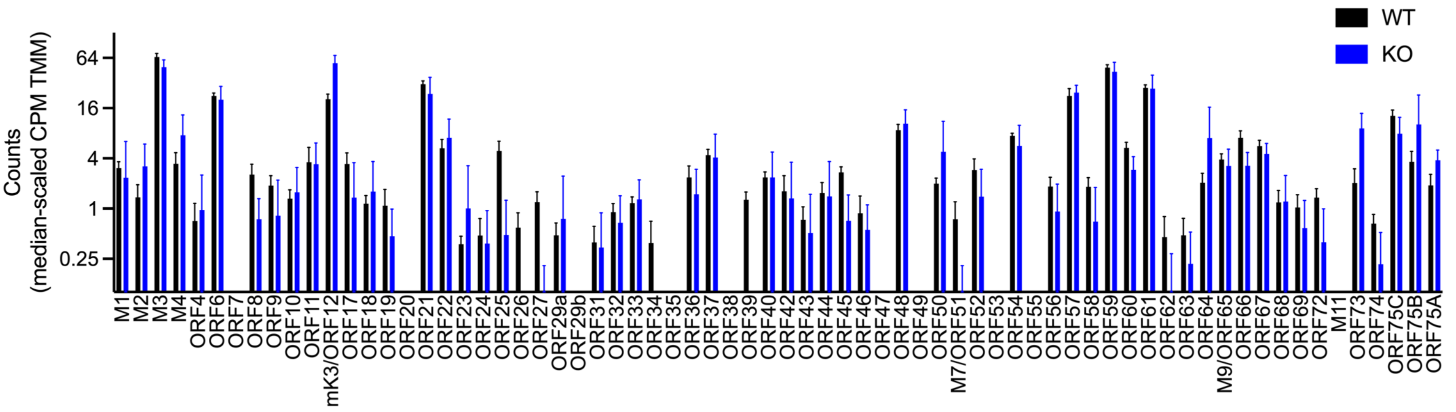

Supplementary Fig. 6

A. Heavy and light chain V-gene usage, mBMC set 1

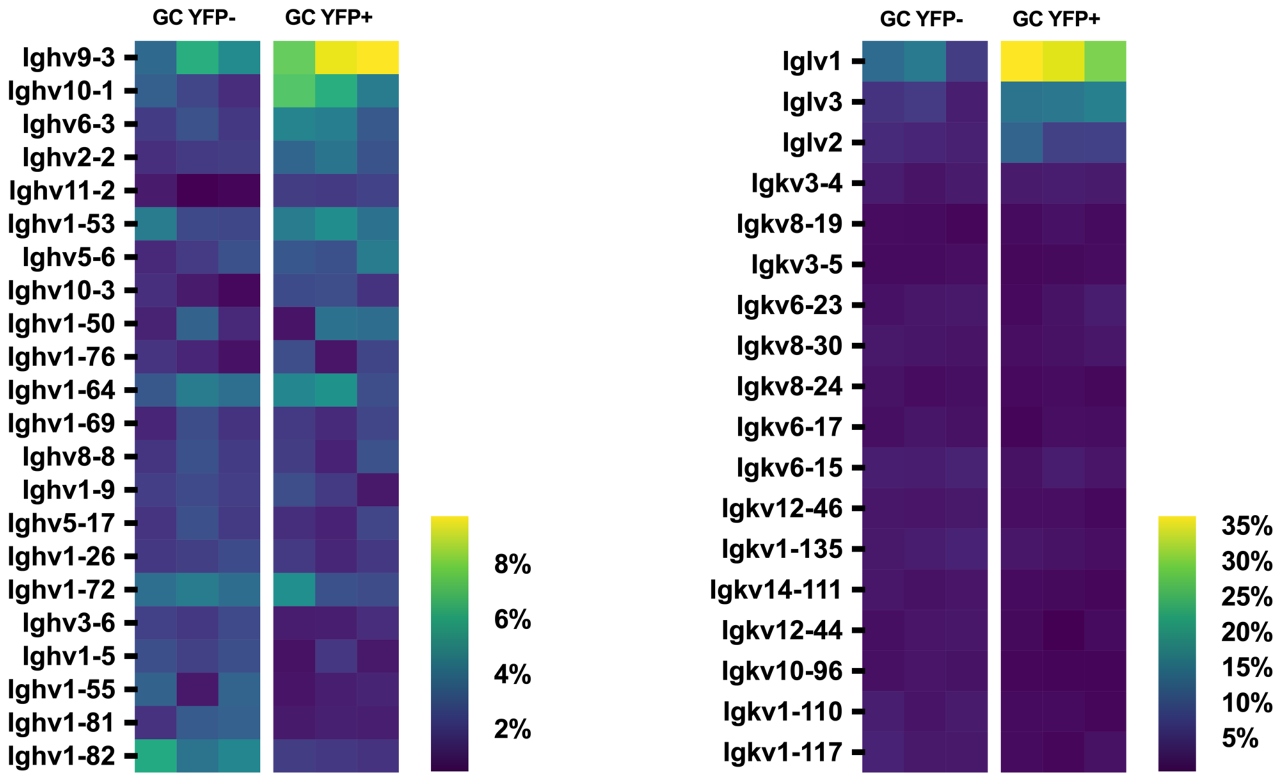


B. Heavy and light chain V-gene usage, mBMC set 2

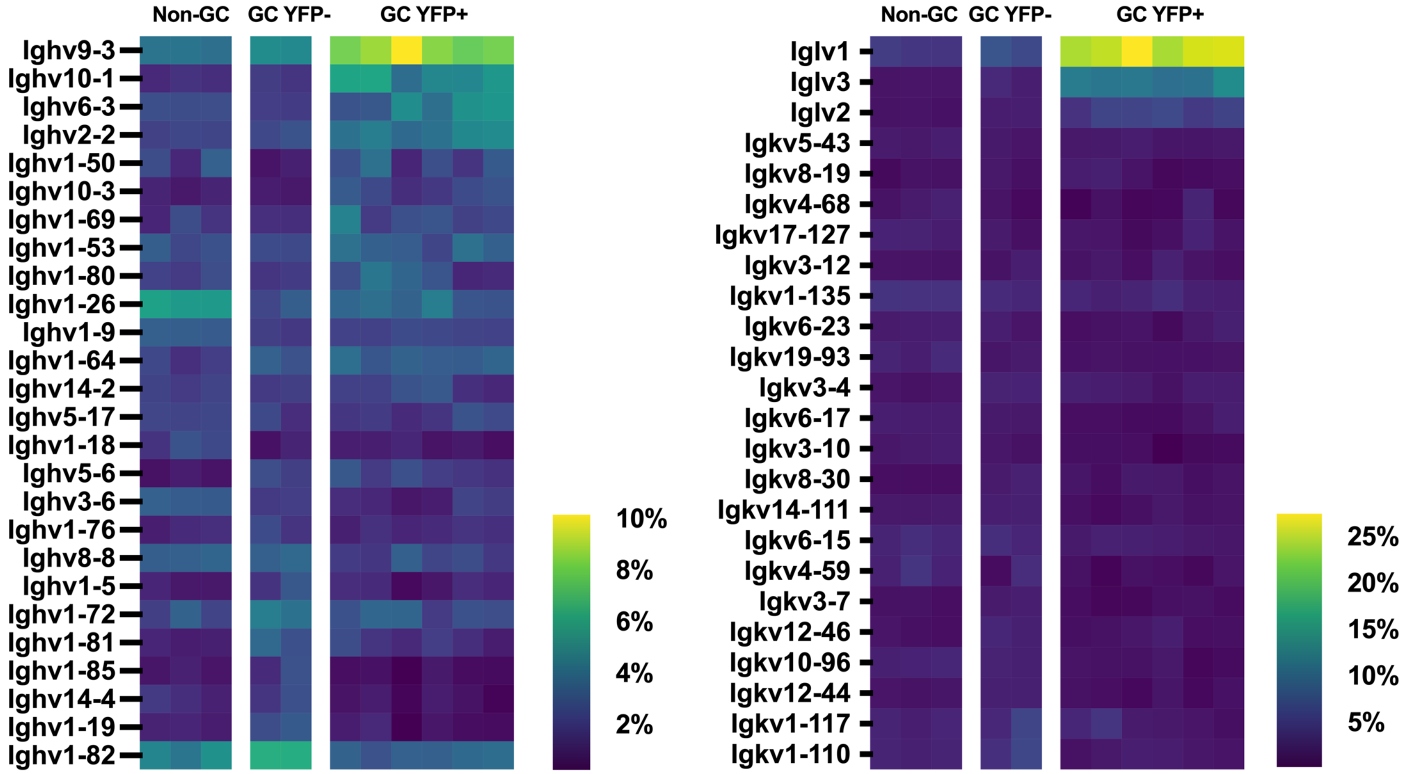

### Supplementary Fig. 7

#### A. GSEA IFN Alpha Response

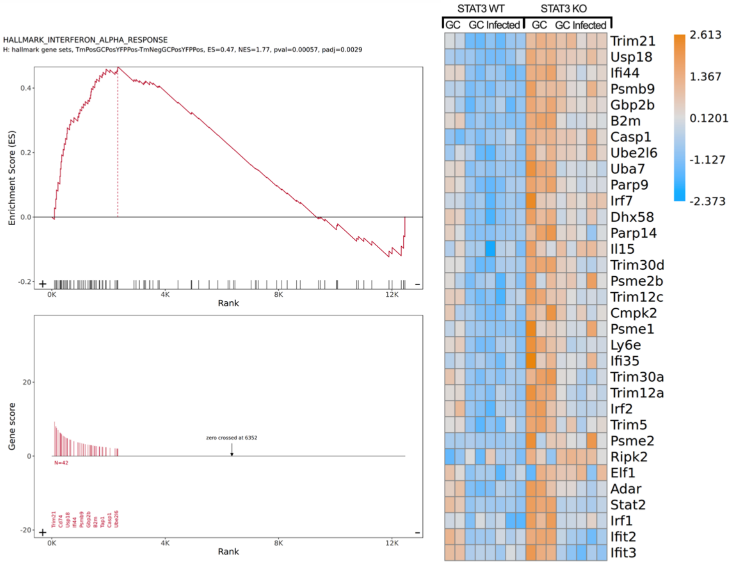


#### B. GSEA IFN Gamma Response

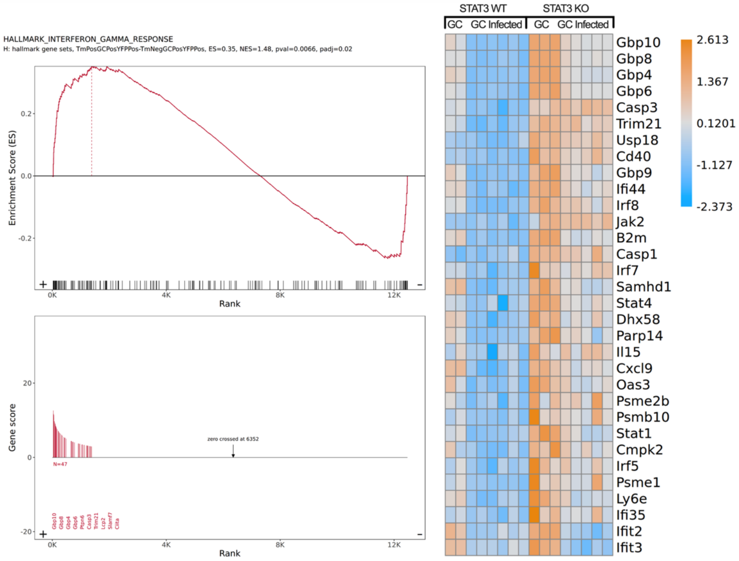


#### C. ISG validation by RT-qPCR

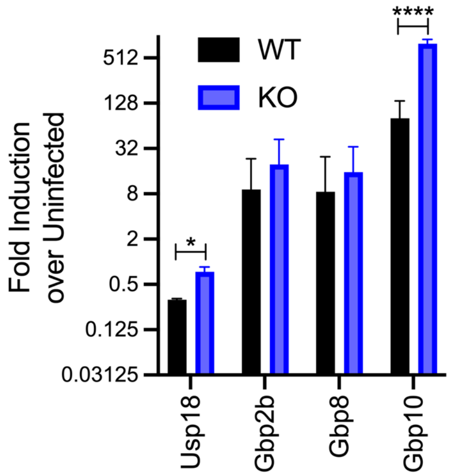
